# Glypican-1 drives unconventional secretion of Fibroblast Growth Factor 2

**DOI:** 10.1101/2021.11.11.468179

**Authors:** Carola Sparn, Eleni Dimou, Annalena Meyer, Roberto Saleppico, Sabine Wegehingel, Matthias Gerstner, Severina Klaus, Helge Ewers, Walter Nickel

## Abstract

Fibroblast Growth Factor 2 (FGF2) is a tumor cell survival factor that is transported into the extracellular space by an unconventional secretory mechanism. Cell surface heparan sulfate proteoglycans are known to play an essential role in this process. Unexpectedly, we found that among the diverse sub-classes consisting of syndecans, perlecans, glypicans and others, Glypican-1 (GPC1) is both the principle and rate-limiting factor that drives unconventional secretion of FGF2. By contrast, we demonstrate GPC1 to be dispensable for FGF2 signaling into cells. We provide first insights into the structural basis for GPC1-dependent FGF2 secretion, identifying disaccharides with N-linked sulfate groups to be enriched in the heparan sulfate chains of GPC1 to which FGF2 binds with high affinity. Our findings have broad implications for the role of GPC1 as a key molecule in tumor progression.

## Introduction

Proteoglycans are components of the extracellular matrix and play essential roles in the storage and protection of growth factors, chemokines and morphogens that bind to the glycosaminoglycan chains of proteoglycans on cell surfaces (Schlessinger et al., 1995; Ori et al., 2011). These post-translational modifications represent unbranched chains of repetitive disaccharide building blocks. They can be classified into four categories defined by (i) heparan sulfates, (ii) chondroitin sulfates, (iii) keratan sulfates and (iv) hyaluronic acid. Heparan sulfates are characterized by about 20-300 negatively charged residues with almost infinite structural modifications such as epimerization and sulfation patterns that are dynamically processed by enzymes resulting in variations between tissues, developmental stages and the type of core protein they are attached to (Turnbull et al., 2001). Different classes of HSPGs also differ in terms of membrane association with glypicans (GPCs) containing a GPI anchor and syndecans (SDCs) having transmembrane spans. This causes lateral segregation of GPCs and SDCs that differentially partition into liquid ordered and disordered domains, respectively, providing a structural basis for distinct roles in growth factor signaling (Gutierrez and Brandan, 2010). As part of their functions to modulate cell growth and differentiation, various kinds of proteoglycans are known to play key roles in tumorigenesis and cancer progression. Their expression patterns along with the structural aspects discussed above and their potential release from cell surfaces play crucial roles in the coordination of their biological functions. These may differ between different kinds of proteoglycans depending on tissue types, developmental stages of tumors and different tumor microenvironments (Blackhall et al., 2001a; De Pasquale and Pavone, 2020b). Proteoglycans are also known to play key roles in the development of chemoresistances making them suitable drug targets for anti-cancer therapies (Lanzi et al., 2017).

Among the proteins that bind to the alternating negatively charged disaccharide units of heparan sulfate chains in HSPGs is FGF2 (Lindahl et al., 1999; Murphy et al., 2004), a pro-angiogenic factor involved in cell proliferation and differentiation during development. In addition, under pathophysiological conditions, FGF2 has a strong impact on tumor-induced angiogenesis triggering the formation of new blood vessels to provide the large demands of malignant cancers for nutrients and oxygen (Carmeliet, 2000; Akl et al., 2015; Akl et al., 2016). FGF2 also plays a critical role as a tumor cell survival factor blocking programmed cell death through both autocrine and paracrine signaling (Okada-Ban et al., 2000; Akl et al., 2016; Traer et al., 2016). Therefore, blocking the biological functions of FGF2 by either limiting its secretion into the extracellular space or inhibiting FGF2 signaling into cells are suitable strategies in anti-cancer treatments (Akl et al., 2016; Pallotta and Nickel, 2020).

While the majority of extracellular proteins contain N-terminal signal peptides for ER-Golgi-dependent protein secretion (Palade, 1975; Rothman, 1994; Rothman and Wieland, 1996; Schekman and Orci, 1996), FGF2 lacks a signal peptide and thus does not have access to the ER/Golgi-dependent secretory pathway (Nickel, 2005; Nickel, 2007). Instead, FGF2 is secreted into the extracellular space by an unconventional mechanism of protein secretion (Rabouille, 2017; Dimou and Nickel, 2018; Pallotta and Nickel, 2020). Various kinds of such pathways have been identified that were collectively termed ‘unconventional protein secretion’ (UPS) (Malhotra, 2013; Rabouille, 2017; Dimou and Nickel, 2018; Pallotta and Nickel, 2020). The mechanism by which FGF2 is transported into the extracellular space is based on direct protein translocation across the plasma membrane (UPS Type I) (Schäfer et al., 2004; Zehe et al., 2006; Rabouille, 2017; Steringer et al., 2017; Dimou and Nickel, 2018; Dimou et al., 2019; Pallotta and Nickel, 2020).

All molecular components known to date to play a role in unconventional secretion of FGF2 are physically associated with the plasma membrane. These factors include the Na,K-ATPase (Zacherl et al., 2015), Tec kinase which is recruited to the inner leaflet via binding to PI(3,4,5)P_3_ (Ebert et al., 2010; Steringer et al., 2012; La Venuta et al., 2016) and PI(4,5)P_2_ (Temmerman et al., 2008; Temmerman and Nickel, 2009; Nickel, 2011), the most abundant phosphoinositide at the inner leaflet of the plasma membrane (Di Paolo and De Camilli, 2006). In the above-mentioned studies, FGF2 has been demonstrated to engage in direct physical interactions with all three of these components with the Na,K-ATPase being the first contact for FGF2 at the inner plasma membrane leaflet (Legrand et al., 2020). Through subsequent interactions with PI(4,5)P_2_ mediated by a cluster of basic amino acids on the molecular surface of FGF2 [K127, R128 and K133;s (Temmerman et al., 2008; Müller et al., 2015; Steringer et al., 2017)], the core mechanism of FGF2 membrane translocation is triggered. This process involves membrane insertion of FGF2 oligomers (Steringer et al., 2012; Steringer et al., 2017; Steringer and Nickel, 2018) whose biogenesis depends on two surface cysteines in FGF2 that drive oligomerization through the formation of intermolecular disulfide bridges (Müller et al., 2015; Steringer et al., 2017; Dimou and Nickel, 2018). Membrane-inserted FGF2 oligomers are accommodated within a lipidic membrane pore with a toroidal architecture (Steringer et al., 2012; Müller et al., 2015; Steringer and Nickel, 2018). This conclusion was derived from several independent observations including simultaneous membrane passage of fluorescent tracers and transbilayer diffusion of membrane lipids triggered by PI(4,5)P_2_-dependent FGF2 oligomerization and membrane insertion (Steringer et al., 2012; Steringer and Nickel, 2018). In further support of this, diacylglycerol, a cone-shaped lipid that interferes with membrane curvature stabilized by PI(4,5)P_2_, was found to inhibit membrane insertion of FGF2 oligomers (Steringer et al., 2012; Steringer and Nickel, 2018), a typical phenomenon for toroidal membrane pores (Gilbert et al., 2014). Based upon these findings, the role of PI(4,5)P_2_ in unconventional secretion of FGF2 has been proposed to be three-fold with (i) mediating recruitment of FGF2 at the plasma membrane, (ii) orienting FGF2 molecules at the inner leaflet to drive oligomerization and (iii) stabilizing local curvature to allow for a toroidal membrane structure surrounding membrane-inserted FGF2 oligomers that are accommodated within a hydrophilic environment (Dimou and Nickel, 2018; Steringer and Nickel, 2018).

As discussed above, membrane-inserted FGF2 oligomers have been proposed to act as key intermediates in FGF2 membrane translocation based on an assembly/disassembly mechanism driving directional transport of FGF2 across the plasma membrane (Dimou and Nickel, 2018; Steringer and Nickel, 2018). This process depends on membrane-proximal heparan sulfate proteoglycans on cell surfaces that capture and disassemble FGF2 translocation intermediates mediating the final step of FGF2 transport into the extracellular space (Zehe et al., 2006; Nickel, 2007; Nickel and Seedorf, 2008; Nickel and Rabouille, 2009). A critical property of heparan sulfates for this function is their ability to out-compete PI(4,5)P_2_ with regard to physical interactions towards FGF2. These are mutually exclusive with heparan sulfates having an about 100-fold higher affinity for FGF2 compared to PI(4,5)P_2_ (Steringer et al., 2017). FGF2 on cell surfaces undergoes intercellular spreading by direct cell-cell contacts, probably mediated by direct exchange between heparan sulfate chains that are physically associated with opposing cell surfaces (Zehe et al., 2006). Thus, during the lifetime of an FGF2 molecule, the role of heparan sulfate proteoglycans is threefold with (i) mediating the final step of FGF2 secretion (Zehe et al., 2006; Nickel, 2007), (ii) protecting FGF2 on cell surfaces against degradation and denaturation (Nugent and Iozzo, 2000) and (iii) mediating FGF2 signaling as part of ternary complexes containing FGF2, heparan sulfate chains and FGF high-affinity receptors (Presta et al., 2005; Ribatti et al., 2007; Belov and Mohammadi, 2013).

In conclusion, based upon sequential interactions of FGF2 with PI(4,5)P_2_ at the inner plasma membrane leaflet and, following the formation of membrane-spanning FGF2 oligomers, interactions with heparan sulfates on cell surfaces, the proposed mechanism of FGF2 membrane translocation offers a molecular basis for directional FGF2 transport into the extracellular space. It has recently been confirmed in a fully reconstituted system using giant unilamellar vesicles (Steringer et al., 2017) and is consistent with earlier observations demonstrating that membrane translocation depends on a fully folded state of FGF2 that permits PI(4,5)P_2_-dependent FGF2 oligomerization and interactions with heparan sulfate chains (Backhaus et al., 2004; Torrado et al., 2009). Furthermore, PI(4,5)P_2_-and heparan-sulfate-dependent translocation of FGF2 across the plasma membrane has also been visualized in living cells using single molecule TIRF microscopy. These studies revealed the real-time kinetics of this process with an average time interval for FGF2 membrane translocation of about 200 ms (Dimou et al., 2019; Pallotta and Nickel, 2020).

In the current study, we made the intriguing and unexpected discovery that HSPGs of different kinds cannot serve equally in capturing FGF2 on cell surfaces as the final step of its unconventional secretory mechanism. Instead, using a proteome-wide BioID screen, we identified GPC1 as the principle HSPG driving this process. Even though HeLa cell lines lacking GPC1 were found to contain normal amounts of total glycosaminoglycans, FGF2 secretion was severely impaired. This phenotype could be reversed by re-expression of GPC1. By contrast, GPC5, the second family member of glypicans expressed in HeLa cells failed to rescue the GPC1 knock-out as did SDC4, an HSPG from the syndecan family. Following the purification of various ectodomains from glypicans and SDC4, the quantification of the binding kinetics revealed a strong preference of FGF2 for GPC1. These findings were corroborated by a strongly increased binding of recombinant FGF2-GFP to the surface of cells overexpressing GPC1. Based on analytical methods, we found disaccharide units enriched in the heparan sulfate chains of GPC1 that are known to play a role in recruiting FGF2. The strong FGF2 binding efficiency of GPC1 could therefore be based on a unique arrangement of these disaccharide units in the minimal pentameric binding motifs in the glycosaminoglycan chains of GPC1. As opposed to its critical role driving efficient secretion of FGF2, we found GPC1 to be dispensable for FGF2 signaling. Our studies reveal a novel und unexpected functional specialization of an HSPG with major implications for the prominent role of GPC1 in tumor progression.

## Results

### GPC1 and FGF2 are in proximity on cell surfaces

To unveil so far unidentified proteins that are in proximity to FGF2 at any time of its lifetime in intact cells, we conducted a proteome-wide BioID screen. A HeLa S3 cell line was generated expressing a fusion protein of FGF2 and the promiscuous biotin ligase BirA (Roux et al., 2012). In a control cell line, a myc-tagged form of BirA was expressed. Both constructs were stably integrated into the genomes of the corresponding HeLa S3 cell lines expressing these fusion proteins in a doxycycline-dependent manner. Following 48 hours of incubation of doxycycline-induced cells in the presence or absence of biotin, a Western analysis was performed to visualize biotinylated proteins under the conditions indicated (Fig. 1A). This analysis revealed distinct patterns of biotinylated proteins when cell lines expressing FGF2-BirA were compared with those expressing BirA alone (Fig. 1A, lane 5 vs. lane 7). By contrast, a post-lysis addition of biotin did not affect the patterns of biotinylated proteins, irrespective of whether conditions with or without biotin in the culture medium were compared (Fig. 1A). These observations indicate that the vast majority of biotinylated proteins was generated in viable cells before lysis.

**Fig. 1:**
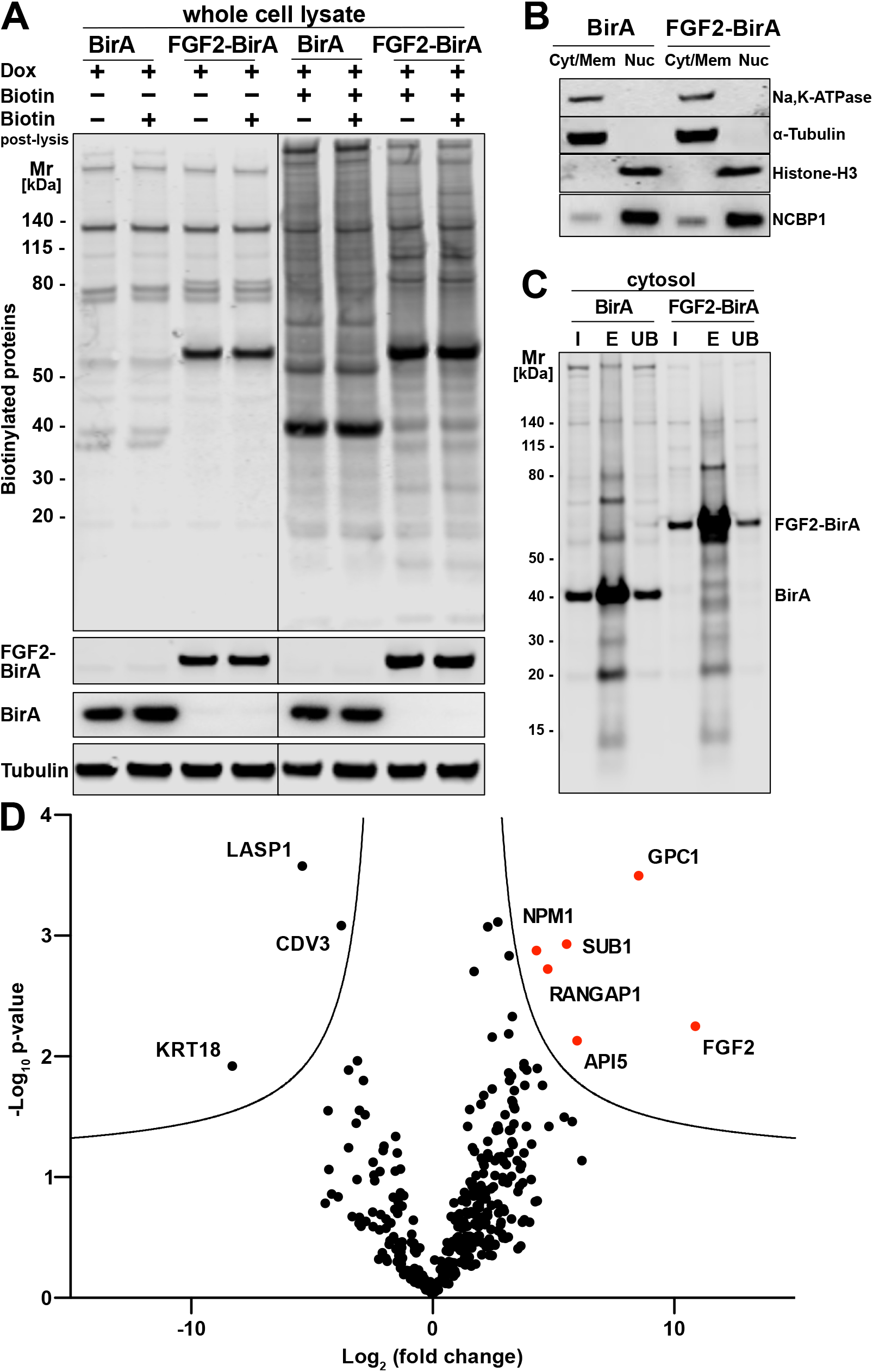
GPC1 and FGF2 are in spatial proximity in a cellular context. (A) HeLa S3 cells stably expressing either FGF2-BirA or myc-tagged BirA (control) in a doxycycline-dependent manner were cultured as detailed in the Materials and Methods section. Whole cell lysates generated from the experimental conditions indicated were subjected to a Western blot analysis. Biotinylated proteins were identified with fluorescent streptavidin. The expression of the fusion proteins was tested with antibodies directed against FGF2 (for FGF2-BirA) or the myc epitope (for BirA). In all samples, tubulin was used as a loading control. (B) HeLa S3 cells expressing FGF2-BirA or myc-tagged BirA were fractionated into nuclei (Nuc) and cellular membranes plus cytosol (Cyt/Mem) as described in the Materials and Methods section. The fractionation was controlled by markers for the plasma membrane (Na,K-ATPase), the cytosol (α-tubulin) and nuclear proteins (Histone-H3 and NCBP1). (C) Large scale preparations of nuclei-free fractions from both FGF2-BirA and myc-tagged BirA expressing cell lines containing biotinylated target proteins. Based on the Cyt/Mem fractionation shown in panel B, all biotinylated proteins were isolated using streptavidin beads. Following elution (lane “E”), all regions except those containing the BirA fusion proteins were extracted and subjected to a quantitative mass spec analysis shown in panel D. For details see Materials and Methods. (D) Biotinylated proteins identified by mass spectrometry and visualized by a Volcano plot indicating hits based on their relative abundance in FGF2-BirA versus myc-tagged BirA-expressing cells. The quantification was based on peptide intensities expressed as “x-fold change” (log_2_; FGF2-BirA/myc-BirA). The experiment was performed in three replicates from which p-values (−log_10_) were calculated (unpaired t-test, two-sided). For further details, see Materials and Methods.

To focus on proteins in proximity of FGF2 that are not localized to the nucleus, a fractionation protocol was established to remove nuclei from all other membranes and cytosolic components. As shown in Fig. 1B and described in detail in Materials and Methods, a fraction containing both α-tubulin (as a cytosolic marker) and the Na,K-ATPase (as a plasma membrane marker) could be generated that is devoid of nuclear markers such as histone-H3 and NCBP1. Based on the procedures described in Figs. 1A and 1B, fractions with nuclear factors being removed were prepared from both FGF2-BirA- and myc-tagged BirA-expressing cells. Biotinylated proteins were pulled down with streptavidin beads, subjected to SDS-PAGE followed by a Western analysis (Fig. 1C). The biotinylated fraction of proteins was subjected to a mass spectrometry analysis to identify all proteins that were in proximity of FGF2 at the level of intact cells (Fingerprints Proteomics Facility at Dundee University, Scotland).

A comparative protein quantification between FGF2-BirA- and myc-BirA-containing fractions was conducted based on peptide intensities. Based on three replicates, for each hit, the differences in peptide intensities between FGF2-BirA and myc-tagged BirA lysates (Log2, fold change) were plotted against the negative Log10 p-value (Fig. 1D). The resulting volcano plot identified proteins in the upper right corner that were more abundant in the FGF2-BirA fraction in a statistically significant manner. This analysis revealed known interaction partners of FGF2 such as API5 (Noh et al., 2014; Bong et al., 2020). In addition, FGF2 itself was identified likely due to its ability to oligomerize at the inner leaflet of the plasma membrane.

The strongest hit of this screen was GPC1, a GPI-anchored heparan sulfate proteoglycan associated with cell surfaces (Fig. 1D). This was a surprising finding for several reasons. First, BioID screens typically return intracellular proteins as hits since ATP is needed to activate biotin to be transferred by BirA to target proteins. Second, no other heparan sulfate proteoglycans were found in proximity of FGF2. These observations were taken as evidence that GPC1 may represent a heparan sulfate proteoglycan that is intimately linked to sites of FGF2 membrane translocation with a specialized function in unconventional secretion of FGF2.

### GPC1 is a rate-limiting component of the FGF2 secretion machinery

Following the identification of GPC1 as a cell surface heparan sulfate proteoglycan in cellular proximity of FGF2 (Fig. 1), we engineered cell lines with knockouts of GPC1 and GPC5, the two family members expressed in HeLa S3 cells (Fig. S1). This included single and double knockouts along with cell lines from each knockout background being stably modified to re-express either GPC1 or GPC5 for rescue experiments (Fig. S1A). Further cell lines were generated expressing each member of the glypican family (GPC1 to 6) in a GPC1 knockout background (Fig. S1B). Finally, we engineered GPC1 knockout cell lines in which a GPC1 version with a transmembrane domain (instead of the natural GPI anchor) or SDC4, a heparan sulfate proteoglycan from the syndecan family (characterized by membrane anchors based on transmembrane spans) were expressed (Fig. S1C).

The engineered HeLa cell lines described in Fig. S1A were analyzed for their ability to secrete FGF2 (Fig. 2). A well-established biotinylation assay was used to quantify FGF2-GFP on cell surfaces [Fig. 2A; (Engling et al., 2002; Seelenmeyer et al., 2005; Zehe et al., 2006; Müller et al., 2015; La Venuta et al., 2016; Legrand et al., 2020)]. A representative Western analysis used for quantification is shown in Fig. 2C. To validate the cell-surface biotinylation experiments, we also quantified FGF2 on cell surfaces using a well-established flow cytometry assay [Fig. 2B; (Engling et al., 2002; Backhaus et al., 2004; Stegmayer et al., 2005; Temmerman et al., 2008; Ebert et al., 2010; Zacherl et al., 2015)]. For both read-outs, all experimental conditions were normalized against HeLa wild-type cells (Fig. 2A and 2B, dotted lines) and differences were evaluated for statistical significance. These experiments revealed a strong decrease in FGF2 secretion efficiency when GPC1 was absent. By contrast, a knockout of GPC5 did not impact this process. Consistently, a double knock-out of GPC1 and GPC5 did not further intensify the FGF2 secretion phenotype observed in cells in which only GPC1 was knocked out. In all cell lines described, overexpression of GPC1 did not only rescue the knock-out of the endogenous GPC1 gene but rather increased the efficiency of FGF2 secretion to levels well above HeLa wild-type cells.

**Fig. 2:**
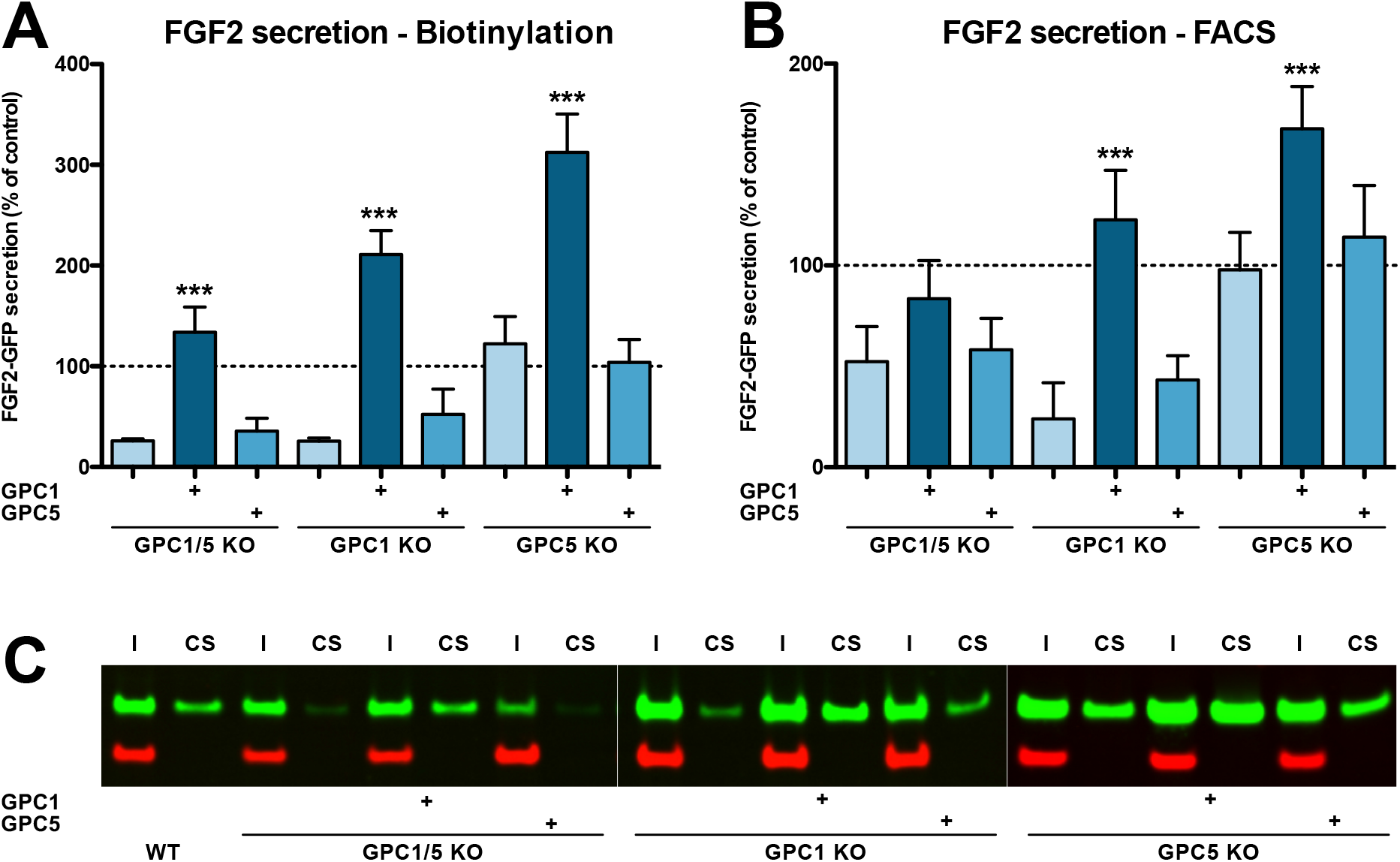
Efficient secretion of FGF2 to cell surfaces depends on GPC1. (A) Quantitative analysis of FGF2 secretion under the experimental conditions indicated measured by cell surface biotinylation. The secretion efficiency of wild-type cells was set to 100%. Standard deviations are shown (n=4). Statistical significance was analyzed using a one-way ANOVA test combined with Tukey’s post hoc test (ns, p > 0.05; *, p ≤ 0.05; **, p ≤ 0.01 and ***, p ≤ 0.001). (B) Quantitative analysis of FGF2 secretion under the experimental conditions indicated measured by analytical flow cytometry. Standard deviations are shown (n=5). Statistical significance was analyzed using a one-way ANOVA test combined with Tukey’s post hoc test (ns, p > 0.05; *, p ≤ 0.05; **, p ≤ 0.01 and ***, p ≤ 0.001). (C) Representative example of a cell surface biotinylation experiment used for the quantitative analysis and statistics shown in panel A (I=input; CS=cell surface) For details, see Materials and Methods.

We further analyzed the cell lines described above with regard to their cell surface capacities to recruit FGF2-GFP (Fig. 3A), their total contents of glycosaminoglycan chains (Fig. 3B) and their total amounts of heparan sulfate chains (Fig. 3C). Using a flow cytometry assay to quantify binding of recombinant FGF2-GFP to cell surfaces, we found that cells lacking GPC1 display slightly reduced binding capacities for FGF2-GFP. By contrast, a GPC5 knock-out did not affect FGF2-GFP binding to cell surfaces. Strikingly, all types of cell lines overexpressing GPC1 were characterized by significantly increased binding capacities for FGF2-GFP. Again, GPC5 overexpression did not result in increased binding of FGF2-GFP to cell surfaces (Fig. 3A). These observations were made despite the fact that there were no significant differences in the total amounts of both glycosaminoglycan chains (Fig. 3B) and heparan sulfate chains (Fig. 3C) in all engineered cell lines analyzed in comparison to HeLa wild-type cells. The combined findings documented in Figs. 2 and 3 suggest that, among the various kinds of heparan sulfate proteoglycans expressed in mammalian cells, GPC1 is the principal component that drives efficient secretion of FGF2. The data further indicate that GPC1 is the rate-limiting factor among the components of the FGF2 secretion machinery that appears to have strong binding capabilities towards FGF2 since GPC1 overexpression causes increased binding of FGF2-GFP to cell surfaces (Fig. 3A) without affecting the total amounts of cellular heparan sulfate chains (Fig. 3C).

**Fig. 3:**
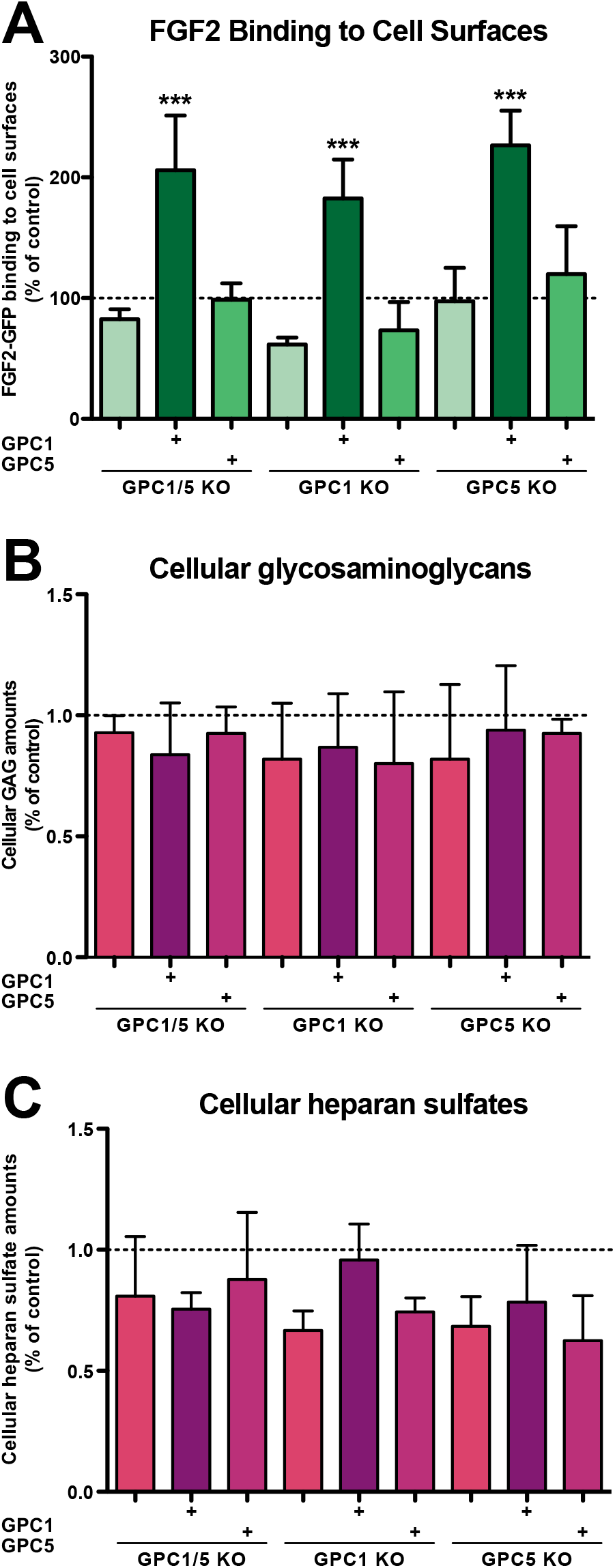
FGF2-GFP binding to cell surfaces is increased in GPC1-overexpressing cells. (A) Quantitative analysis of the FGF2-GFP binding capacity of cell surfaces under the experimental conditions indicated using flow cytometry. Standard deviations are shown (n=5). Statistical significance was analyzed using a one-way ANOVA test combined with Tukey’s post hoc test (ns, p > 0.05; *, p ≤ 0.05; **, p ≤ 0.01 and ***, p ≤ 0.001). (B) Quantification of the total amounts of GAG chains under the experimental conditions indicated. Standard deviations are shown (n=4). Statistical significance was analyzed using a one-way ANOVA test combined with Tukey’s post hoc test (ns, p > 0.05; *, p ≤ 0.05; **, p ≤ 0.01 and ***, p ≤ 0.001). (C) Quantification of the total amounts of heparan sulfate chains under the experimental conditions indicated. Standard deviations are shown (n=3). Statistical significance was analyzed using a one-way ANOVA test combined with Tukey’s post hoc test (ns, p > 0.05; *, p ≤ 0.05; **, p ≤ 0.01 and ***, p ≤ 0.001). For details, see Materials and Methods.

Based on the cell lines described and characterized in Figs. S1, 2 and 3, we tested whether overexpression of other GPC family members can rescue FGF2 secretion in the context of a GPC1 knock-out (Fig. 4). Of note, the GPC family can be divided into two subclasses, GPC1/2/4/6 and GPC3/5. As shown in Fig. 4A and 4C, like GPC5, GPC3 was incapable of promoting efficient FGF2 secretion in the absence of GPC1. By contrast, all GPCs belonging to the GPC1 subfamily rescued FGF2 secretion in a GPC1 knock-out background. While GPC2 and GPC4 did so at the level of HeLa wild-type cells, GPC6 overexpression increased the efficiency of FGF2 secretion above HeLa wild-type levels. Nevertheless, GPC1 overexpression was found to represent the strongest stimulator of FGF2 secretion. As shown in Fig. 4B, these findings were closely reflecting the ability of the various GPC family members to increase the cell surface binding capacities for FGF2-GFP.

**Fig. 4:**
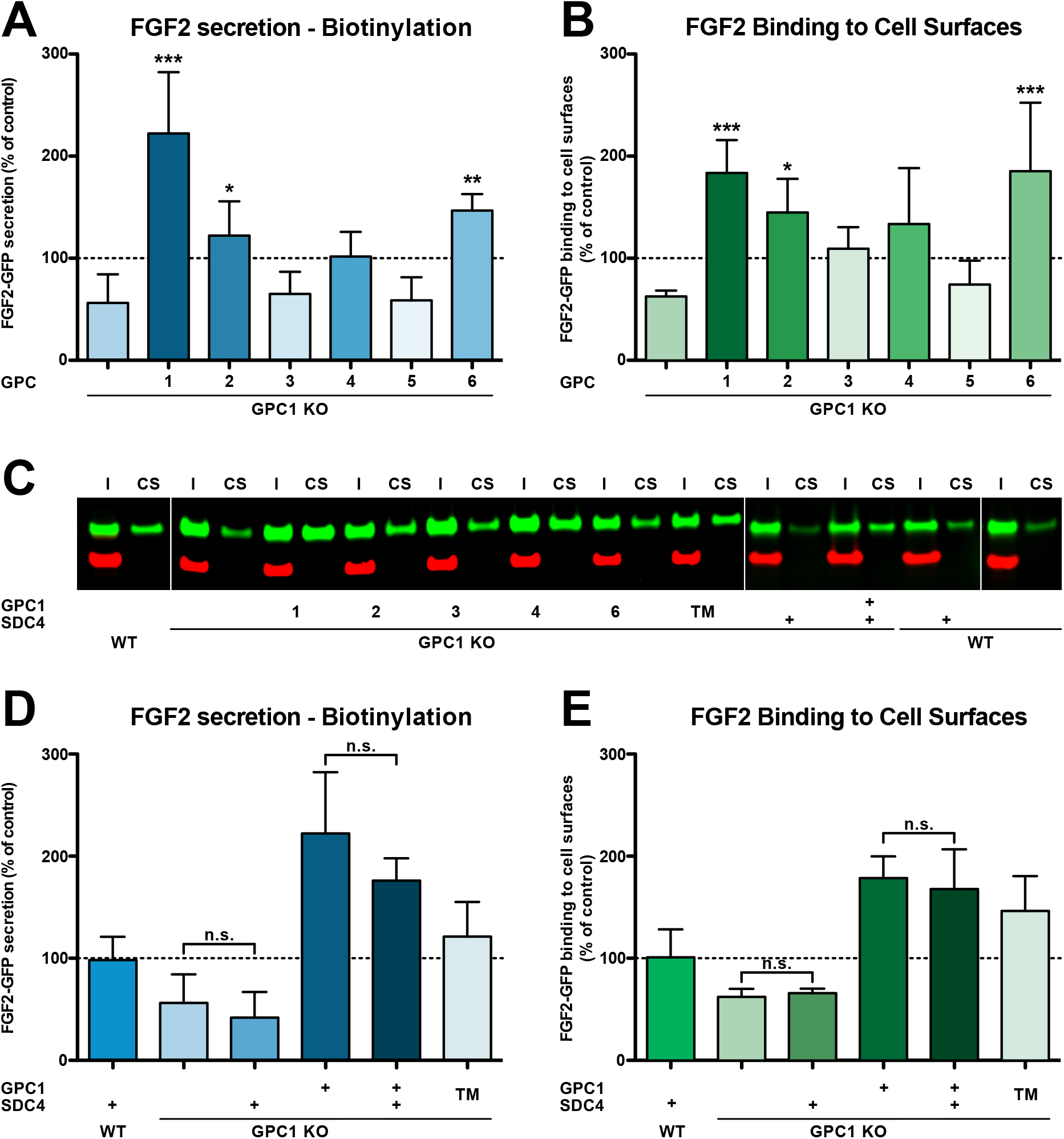
GPC1 is the principal heparan sulfate proteoglycan involved in unconventional secretion of FGF2. (A) Quantitative comparison of all six glypican family members with regard to their potential to drive FGF2 secretion upon overexpression in a GPC1 knockout background based on cell surface biotinylation experiments. Standard deviations are shown (n=5). (B) Quantitative comparison of all six glypican family members with regard to their ability to affect the cell surface binding capacities for FGF2-GFP. The GPCs indicated were overexpressed in a GPC1 knockout background. FGF2-GFP binding to cell surfaces was analyzed by flow cytometry. Standard deviations are shown (n=4). (C) Representative example for the raw data of cell surface biotinylation experiments used to quantify and to statistically evaluate unconventional secretion of FGF2 under the conditions indicated in panel A and D (I=input; CS=cell surface). (D) Quantitative comparison of GPC1 versus SDC4 (syndecan 4) with regard to their potential to drive FGF2 secretion upon overexpression in a GPC1 knockout background based on cell surface biotinylation experiments. “TM” stands for a GPC1 construct in which the GPI anchor was replaced by the membrane span of SDC4. Standard deviations are shown (n=5). (E) Quantitative comparison of GPC1 versus SDC4 with regard to their ability to affect the cell surface binding capacities for FGF2-GFP. FGF2-GFP binding to cell surfaces was analyzed under the experimental conditions indicated using flow cytometry. “TM” stands for a GPC1 construct in which the GPI anchor was replaced by the membrane span of SDC4. Standard deviations are shown (n=4). All statistical analyses were based on a one-way ANOVA combined with Tukey’s post hoc test (ns, p > 0.05; *, p ≤ 0.05; **, p ≤ 0.01 and ***, p ≤ 0.001).

We also tested whether other heparan sulfate proteoglycans such as syndecans can support efficient secretion of FGF2 in a GPC1 knock-out background. As shown in Figs. 4D and 4E, SDC4 overexpression neither rescued FGF2 secretion nor did it affect the cell surface binding capacities for recombinant FGF2-GFP. This observation was not due to the fact that SDC4 and GPC1 structurally differ in membrane attachment with SDC4 carrying a transmembrane span and GPC1 having a GPI anchor. This was particularly evident from the fact that an engineered version of GPC1 in which the GPI anchor was replaced by a membrane span (“GPC1 TM”) was fully functional in GPC1 knock-out cells, both with regard to efficient secretion of FGF2 (Fig. 4D) and cell surface binding capacities for recombinant FGF2-GFP (Fig. 4E).

To assess the impact of GPC1 overexpression on the efficiency by which FGF2 is secreted from cells at different FGF2-GFP expression levels, we used an advanced TIRF assay with single molecule resolution (Dimou et al., 2019). These experiments were conducted in CHO cells that express FGF2-GFP in a doxycycline-dependent manner, reading out the number of FGF2-GFP particles on the cell surface of individual cells (Fig.5). Two conditions were chosen characterized by high (Fig. 5A) and low (Fig. 5C) expression levels of FGF2-GFP. When cells were analyzed expressing FGF2-GFP at high levels, the average number of FGF2-GFP particles on cell surfaces was increased by about 50% in a pool of GPC1 overexpressing cells relative to CHO wild-type cells (Fig. 5B). This difference was even more pronounced in cells expressing low levels of FGF2-GFP resulting in a more than 4-fold higher average number of FGF2-GFP particles on the cell surfaces of GPC1-overexpressing cells compared to wild-type cells (Fig. 5D). Since the pool of GPC1 overexpressing cells was characterized by a range of GPC1 expression levels, a certain heterogeneity of this effect was observed. Strikingly, individual cells were observed that were characterized by a more than 20-fold increase of FGF2-GFP particles on their cell surface compared to the average number of FGF2-GFP particles on the cell surfaces of wild-type cells (Fig. 5D). These findings suggest that GPC1 has an even stronger impact on this process when the amounts of FGF2-GFP being expressed are limiting.

**Fig. 5:**
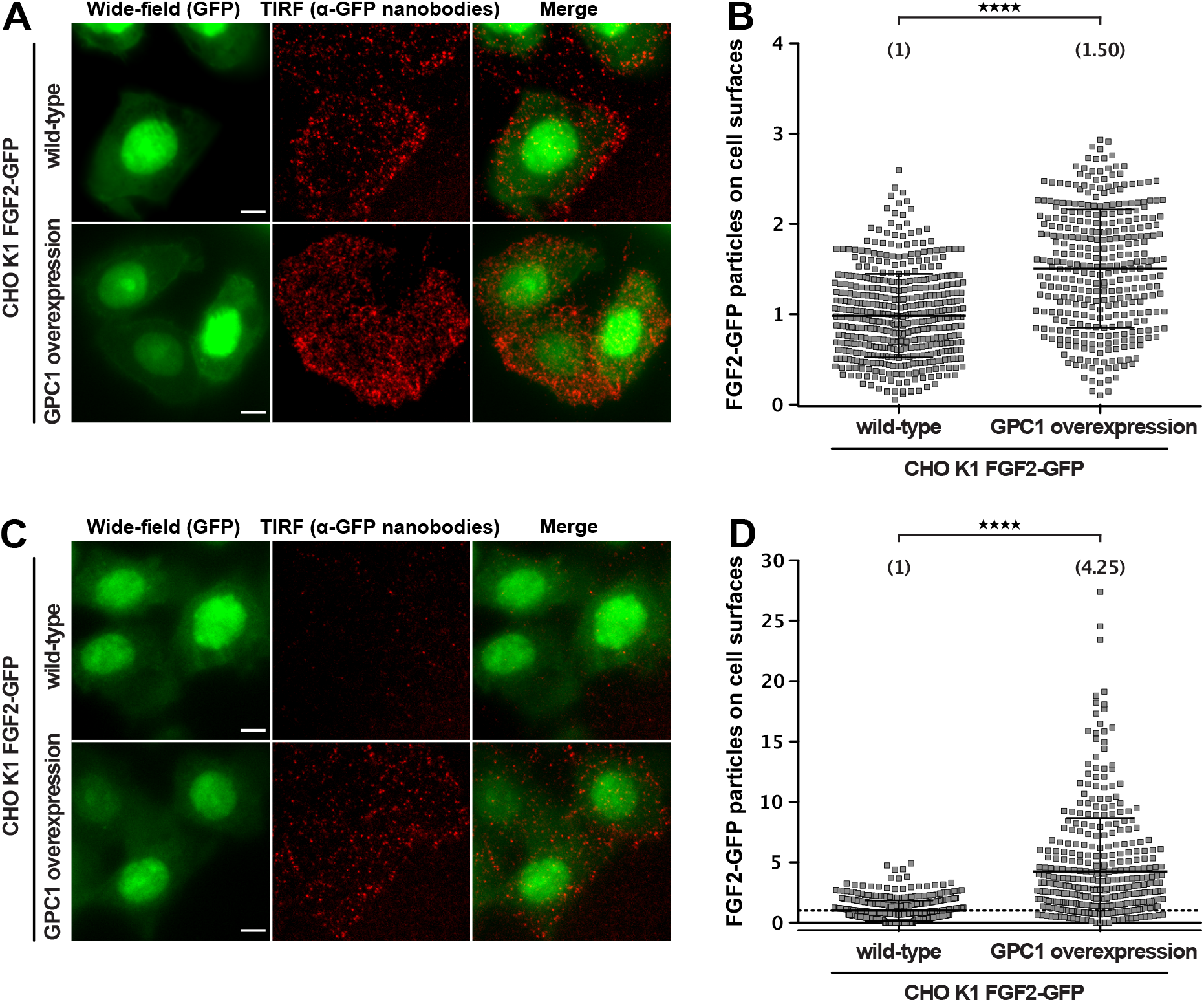
GPC1 overexpression results in increased FGF2 secretion efficiencies. FGF2 secretion efficiencies in wild-type and GPC1 overexpressing cells were assessed by TIRF microscopy using anti-GFP nanobodies to detect single FGF2-GFP molecules on cell surfaces as described earlier (Dimou et al., 2019). For details, see Materials and Methods. (A) Representative examples under experimental conditions at high FGF2-GFP expression levels (B) Quantification and statistical analysis of experiments corresponding to the experimental conditions shown in panel A (C) Representative examples under experimental conditions at low FGF2-GFP expression levels. EM Gain of the wide-field (GFP) was increased for this condition, in order to allow selection of cell area for subsequent quantification (D) Quantification and statistical analysis of experiments corresponding to the experimental conditions shown in panel B Data are shown as mean ± SD (n = 4) (panel B and D). The secretion efficiency of the wild-type cells was set to 1; in panel D, a dotted line was put at 1, to facilitate visualization. The statistical analysis was based on an unpaired t-test (****, p < 0.0001).

The combined findings shown in Figs. 2, 3, 4 and 5 provide strong evidence that GPC1 is the principal heparan sulfate proteoglycan that drives the unconventionally secretory mechanism by which FGF2 is transported to the extracellular surfaces of cells.

### GPC1 and FGF2 form a strong pair of interaction partners

To study the interaction between the heparan sulfate chains from various kinds of glypicans and syndecans with FGF2 at the molecular level, we generated constructs of GPC1, GPC5, GPC6 and SDC4 to express and purify soluble variants of them from mammalian HEK cells (Fig. S2) (Svensson et al., 2009). In case of GPC1, to be used as a negative control, an additional variant form was generated that is defective regarding the addition of heparan sulfate chains (GPC1-ΔHS). As shown in Fig. S2, all purified glypican family members were treated with heparinase III to demonstrate the presence of O-linked heparan sulfate chains. For SDC4, treatments with both heparinase III and chondroitin sulfate degrading enzyme were conducted to reveal the different types of O-glycosylation of this type of proteoglycan. In addition, recombinant FGF2 was purified to homogeneity (Fig. S2).

To study physical interactions of FGF2 with the O-linked heparan sulfate chains of the proteoglycans indicated, we chose biolayer interferometry as read-out (Fig. 6). Following biotinylation, all proteoglycans indicated were immobilized on BLI sensors. The sensors were then brought into contact with a range of FGF2 concentrations between 0.8 and 60 nM. This approach allowed for a quantitative comparison of the binding preferences of FGF2 to various kinds of heparan sulfate chains linked to different types of proteoglycans. It revealed a strong interaction of FGF2 with GPC1 that was detectable already at the lowest FGF2 concentration being used at 0.8 nM (Fig. 6A). By contrast, even at the highest concentration of FGF2 (60 nM), GPC5 only showed weak interactions with FGF2 (Fig. 5C) that were barely above the levels of the negative controls, GPC1-ΔHS (Fig. 6B) or heparan sulfate binding mutants of FGF2 tested against GPC1 (Fig. S3). Unlike GPC5, GPC6, a member of the GPC1 sub-family of glypicans that was capable of rescuing a GPC1 knockout (Fig. 4A and 4B) displayed significant binding capabilities of FGF2, however, less efficiently compared to GPC1 (Fig. 6D). Finally, similar to GPC5, SDC4, a member of the syndecan family of proteoglycans, showed weak interactions with FGF2 at concentrations of up to 60 nM (Fig. 6E).

**Fig. 6:**
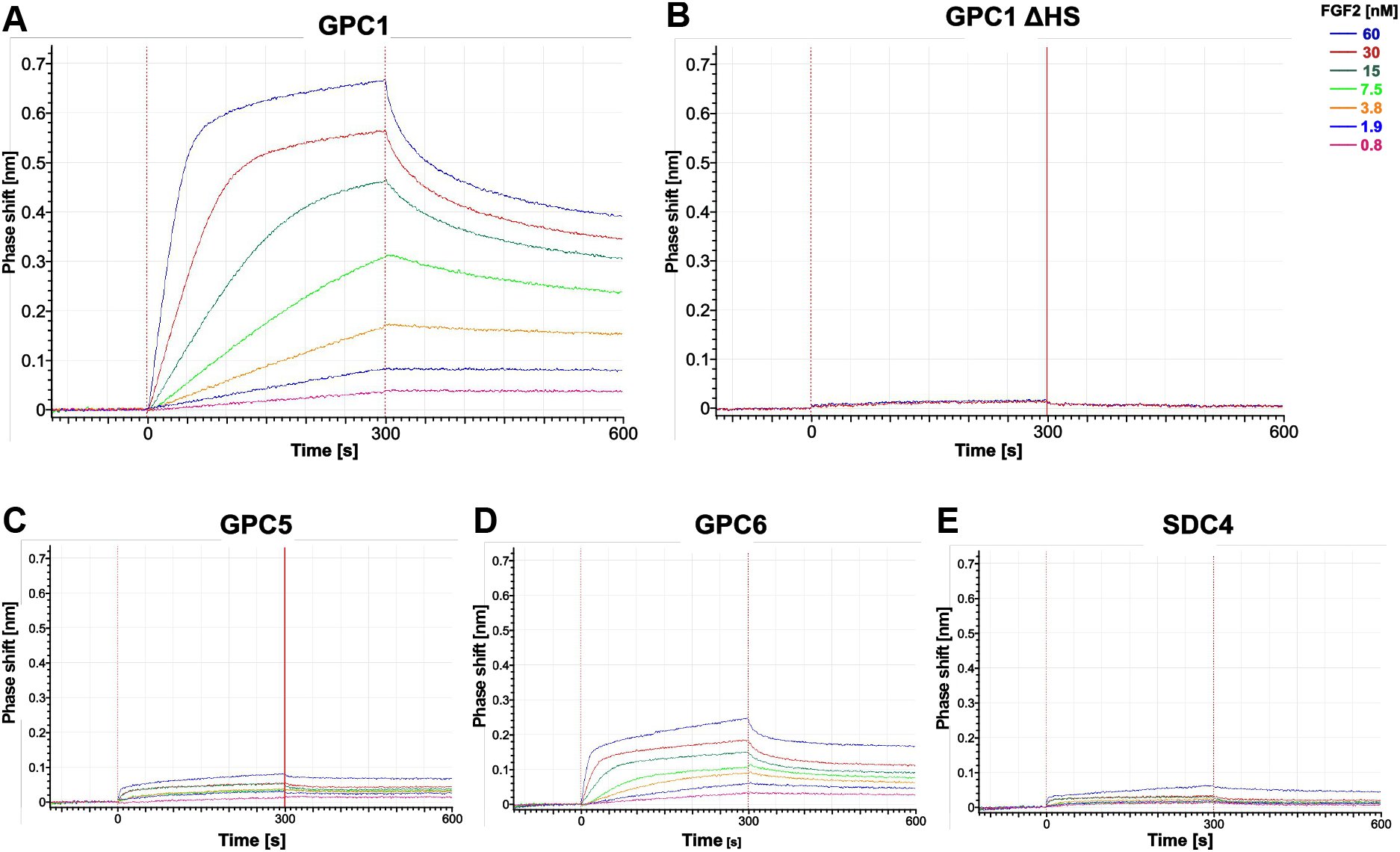
GPC1 and FGF2 form a strong pair of interaction partners. Recombinant constructs encoding soluble ectodomains of GPC1 (panel A), GPC1 ΔHS (panel B; a mutant form to which heparan sulfate chains cannot be added), GPC5 (panel C), GPC6 (panel D) and SDC4 (panel E; a member of the syndecan family of heparan sulfate proteoglycans) were expressed and purified from HEK293 cells (see Fig. S2). Using biolayer interferometry, interactions studies with temporal resolution visualizing both association and dissociation kinetics were conducted with purified FGF2 (Fig. S2) at the concentrations indicated. The data shown are representative for two independent experiments. Experimental details are given in the Materials and Methods section.

As documented in Fig. S3, beyond further controls using heparan sulfate binding mutants of FGF2, we found GPC1 to exhibit only weak or no interactions with other examples of growth factors or cytokines such as EGF and IFNγ. Similarly, examples for other extracellular proteins secreted by unconventional means such as galectin-1 and galectin-3 (Rabouille, 2017; Dimou and Nickel, 2018; Pallotta and Nickel, 2020) were observed not to be capable of interacting with GPC1 (Fig. S3).

The combined findings shown in Figs. 6, S2 and S3 reveal a tight relationship between GPC1 and FGF2 that form a strong pair of interaction partners compared to other proteoglycans, growth factor and cytokines including examples of other proteins secreted by unconventional means. They are consistent with the prominent role of GPC1 as the driver of the unconventional secretory mechanism of FGF2 as shown in Figs 2–5.

### The heparan sulfate chains of GPC1 are enriched in disaccharides known to be critical for FGF2 recruitment

To obtain insight into the molecular mechanism underlying the strong interaction between GPC1 and FGF2, we aimed at analyzing the disaccharide contents of the heparan sulfate chains of GPC1 in comparison to GPC5 and SDC4. Since there is no methodology available to sequence the disaccharide units of heparan sulfate chains, we treated the recombinant purified forms of GPC1, GPC5 and SDC4 (Fig. S2) with a mixture of heparinase I, II and III to convert their heparan sulfate chains into disaccharides. Using an established HPLC protocol [(Carnachan and Hinkley, 2017); see Materials and Methods for details], a total of 12 different heparan sulfate disaccharide standards with different sugar combinations and sulfation patterns (Iduron, UK) were analyzed for their retention times on a HPLC ion exchange column (Fig. S4). These were compared with the retention times of the spectrum of dissacharide units released from GPC1, GPC5 and SDC4 upon treatments with heparinases (Fig. 7A). The relative abundances of each of the identified disaccharides in the heparans sulfate chains of GPC1, GPC5 and SDC4, respectively, were quantified. The observed differences were tested for statistical significance (Fig. 7B). This analysis revealed the enrichment of disaccharides in GPC1 over GPC5 that correspond to the disaccharide standards 1, 2 and 5. All of these disaccharides contain N-linked sulfates (Fig. S4). The biggest difference between GPC1 and GPC5 was found to be the disaccharide standard 1 that represents a tri-sulfated disaccharide with two O-linked and one N-linked sulfate group. Intriguingly, when FGF2 was co-crystalized with synthetic heparin molecules, the binding site was found to contain three disaccharides of the type represented by the disaccharide standard 1 (Raman et al., 2003). By contrast, disaccharides lacking both O- and N-linked sulfations corresponding to the disaccharide standards 6 and 12 (Fig. S4) were more abundant in GPC5 compared to GPC1 (Fig. 7B). When the spectrum of disaccharides from GPC1 and GPC5 was compared with SDC4, most features were similar to GPC1 (disaccharide standards 1, 2, 6 and 12) while the abundance of the disaccharide corresponding to standard 5 was rather similar to GPC5 (Fig. 7A and 7B). These findings suggest that, beyond the overall abundance of certain sulfated disaccharide units in heparan sulfate chains, their combination into trimers of sulfated disaccharides plays a key role in forming a high affinity binding site for FGF2. The strong interaction of GPC1 with FGF2 (Fig. 6) therefore indicates that GPC1 carries heparan sulfate units consisting of three disaccharides corresponding to standard 1 (Fig. S4) as identified in structural in vitro studies (Raman et al., 2003). Based on our findings, such binding sites are likely to be less abundant in GPC5 and SDC4 resulting in weaker interactions with FGF2 (Fig. 6). This, in turn, explains the predominant function of GPC1 in unconventional secretion of FGF2 as demonstrated in Figs 2–5.

**Fig. 7:**
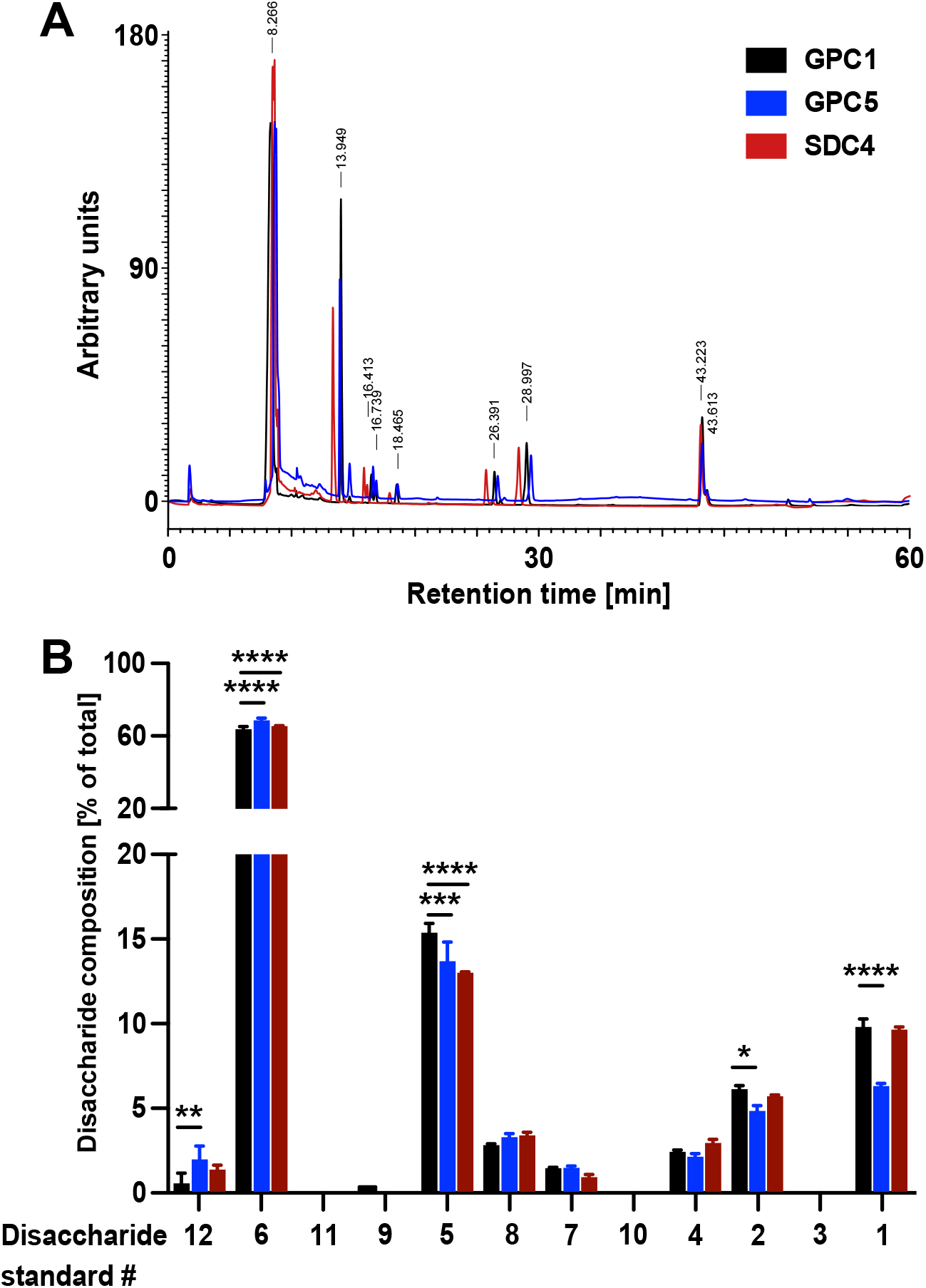
Quantitative characterization of the disaccharide contents of heparan sulfate chains derived from GPC1, GPC5 and SDC4. The recombinant forms of GPC1, GPC5 and SDC4 described in Figs. 6 and S2 were treated with a mixture of the heparinases I + II and III to release the disaccharide units of their heparan sulfate chains. These were then identified by their retention times in a analytical anion exchange HPLC setup using synthetic disaccharides as standards (Fig. S4). For details, see Materials and Methods. (A) Representative elution profile of the disaccharide units derived from the heparan sulfate chains of GPC1 (black), GPC5 (blue) and SDC4 (red) (B) Statistical analysis of four independent experiments providing the relative abundances of heparan sulfate disaccharide units corresponding to the twelve standards (Fig. S4) contained in GPC1 (black), GPC5 (blue) and SDC4 (red). Statistics were based on a two-way ANOVA test combined with a Bonferroni post-test with ns, p > 0.05; *, p ≤ 0.05; **, p ≤ 0.01; ***, p ≤ 0.001 and ****, p ≤ 0.0001.

### GPC1 is dispensable for FGF2-induced ERK1/2 signaling

To test as to whether GPC1 is not only the key driver of FGF2 secretion but also plays a role in FGF2 signal transduction, we analyzed the ability of recombinant FGF2 to initiate signal transduction in GPC1 knockout versus wild-type versus GPC1-overexpressing cells (Fig. 8). As a read-out, we quantified ERK1/2 phosphorylation, an event that occurs downstream of FGF receptor activation. As a positive control, we treated the different cell types indicated with heparinases I, II and III to degrade cell surface heparan sulfates down to their disaccharide subunits. The latter are incapable of forming ternary FGF signaling complexes on the surfaces of target cells consisting of FGF2, high affinity FGF receptors and heparan sulfate chains. At both 10 and 1 ng/ml FGF2, all types of cells indicated showed reduced FGF2 signaling when treated with heparinases [Fig. 8A (quantification with statistics) and 8B (representative Western analysis of ERK phosphorylation)]. By contrast, neither a knock-out nor overexpression of GPC1 had any significant impact on phosphorylated ERK1/2 levels at both 10 and 1 ng/ml FGF2 added to cells (Fig. 8A and 8B). These findings reveal a differential role of GPC1 in FGF2 related processes with GPC1 being essential for efficient secretion of FGF2 but being dispensable for the transmission of FGF2-dependent signals into cells.

**Fig. 8:**
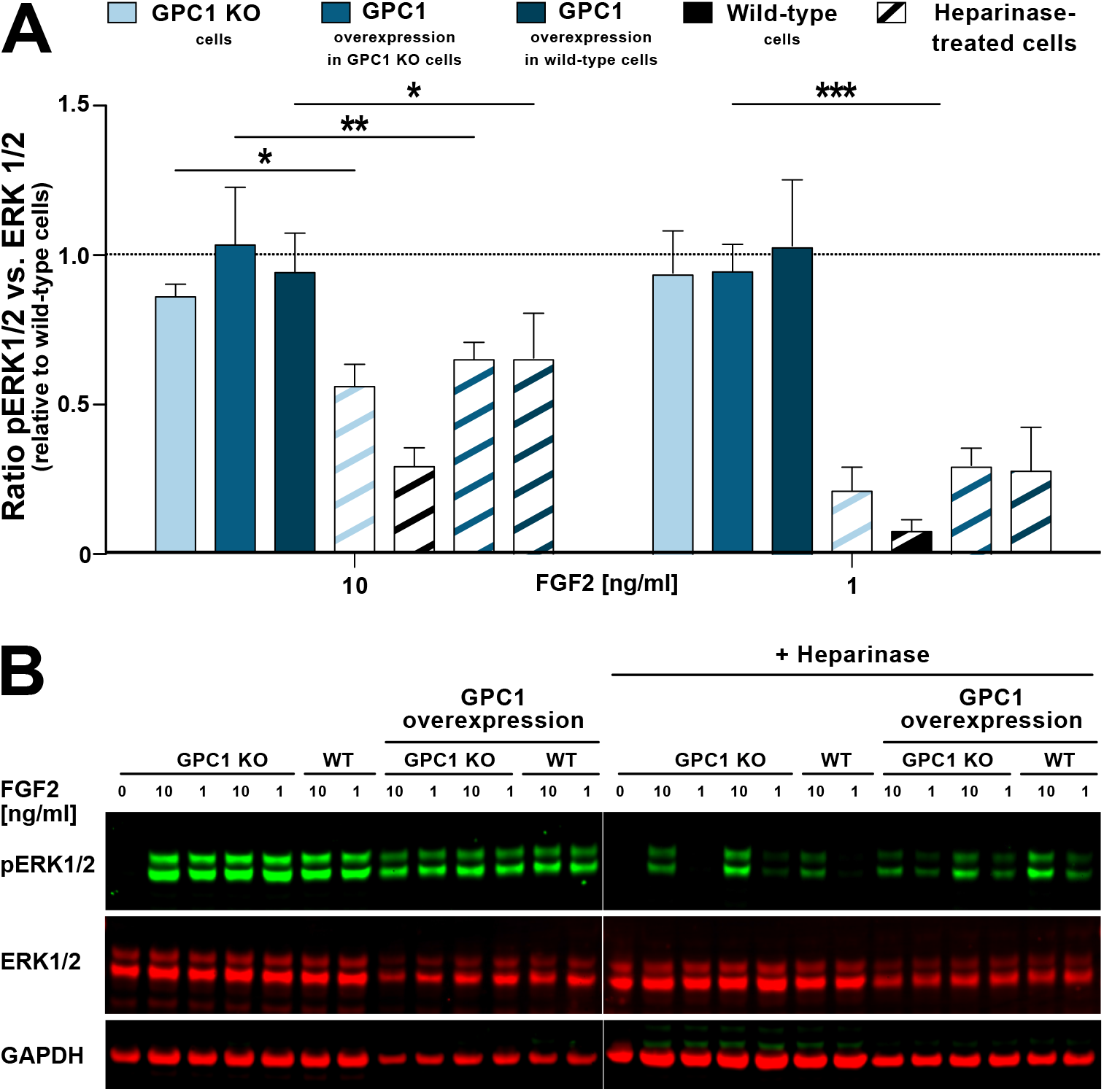
GPC1 is dispensable for FGF2 signaling. Various forms of HeLa cells including wild-type and GPC1 knock-out cells as well as cells overexpressing GPC1 in either a wild-type or a GPC1 knockout background were treated with recombinant FGF2 at the concentrations indicated. Where indicated, cells were treated with a mixture of heparinase I, II and III used as a positive control. As a read-out for FGF signaling, the ratio between phosphorylated and unphosphorylated ERK1/2 was determined. For experimental details, see Materials and Methods. (A) Quantitative analysis of the pERK1/2 to ERK1/2 ratio (n=4; standard deviations are shown). The statistical analysis was based on a one-way ANOVA test combined with Tukey’s post hoc test (ns, p > 0.05; *, p ≤ 0.05; **, p ≤ 0.01 and ***, p ≤ 0.001). (B) A representative Western analysis was used for the quantification and statistical analysis shown in panel A. The GAPDH signal was used as a loading control.

## Discussion

In this study, we report on the surprising identification of the glypican family member GPC1 as a heparan sulfate proteoglycan with a specialized function in driving unconventional secretion of FGF2. While we found that the total amounts of glycosaminoglycans including heparan sulfate chains are not significantly altered in GPC1 knockout cells, we observed a pronounced decrease in FGF2 secretion efficiencies in the absence of GPC1. Overexpression of GPC5 (the second glypican family member expressed alongside GPC1 in HeLa cells) or SDC4, a member of the syndecan family of heparan sulfate proteoglycans, did not rescue this process in a GPC1 knockout background. By contrast, overexpression of GPC1 in GPC1 knockout cells not only restored but rather caused a substantial increase of FGF2 secretion efficiencies. Therefore, GPC1 is a rate-limiting component of the FGF2 secretion machinery that is required for efficient transport of FGF2 into the extracellular space. These findings have implications for the molecular mechanism by which GPC1 is functioning in this process. Beyond the role of cell surface heparan sulfate chains in capturing and disassembling FGF2 oligomers at the outer plasma membrane leaflet (Zehe et al., 2006; Nickel, 2007; Rabouille, 2017; Dimou and Nickel, 2018; Pallotta and Nickel, 2020), they further suggest that GPC1 is already required for membrane insertion of FGF2 oligomers, an upstream step that is initiated at the inner plasma membrane leaflet. Based upon the unique positioning of the heparan sulfate chains of GPC1 in close proximity to its GPI anchor and therefore the membrane surface (Prydz and Dalen, 2000; Blackhall et al., 2001b; Nakato and Kimata, 2002; De Pasquale and Pavone, 2020a), the FGF2 binding sites in GPC1 appear to be required for stabilizing the first subunits of membrane-spanning FGF2 oligomers as they surface at the outer plasma membrane leaflet. In this way, the likeliness of these intermediates to be formed could be strongly increased when GPC1 is overexpressed. This, in turn, would result in FGF2 membrane translocation events occurring more frequently. These observations are consistent with previous findings demonstrating that, compared to wild-type cells, higher forms of membrane-inserted FGF2 oligomers do not accumulate in cells lacking cell surface heparan sulfates (Dimou et al., 2019).

To obtain insight into the molecular mechanism by which GPC1 drives unconventional secretion of FGF2, we conducted quantitative binding studies of FGF2 with GPC1 and other heparan sulfates proteoglycans using recombinant components purified from HEK cells. These experiments revealed GPC1 to be the strongest FGF2 binding partner with the heparan sulfate chains being essential for this interaction. Based on a quantitative analysis of the disaccharide subunits in the heparan sulfate chains of GPC1, GPC5 and SDC4, we found disaccharides with N-linked sulfate groups corresponding to the standards 1, 2 and 5 enriched in GPC1 over GPC5 and SDC4. In particular, a tri-sulfated disaccharide (corresponding to standard 1 in Figs. 7 and S4) was found enriched in GPC1 over GPC5. By contrast, disaccharides lacking O- or N-linked sulfates (such as the ones corresponding to standards 6 and 12) were found enriched in GPC5 and SDC4 compared to GPC1. Consistently, a trimer of the tri-sulfated disaccharide enriched in GPC1 (corresponding to standard 1) has been found to be a strong binding motif for FGF2 in structural in vitro studies using synthetic heparin molecules (Raman et al., 2003). While it remains a goal for future studies to identify the precise FGF2 binding site and its frequency of occurrence in the heparan sulfate chains of GPC1, our findings provide a plausible explanation for the observed differences regarding the FGF2 binding efficiencies towards GPC1, GPC5 and SDC4. Along with the unique spatial organization of the heparan sulfate chains of GPC1 being arranged in close proximity to the plasma membrane surface (Prydz and Dalen, 2000; Blackhall et al., 2001b; Nakato and Kimata, 2002; De Pasquale and Pavone, 2020a), the identification of GPC1 being a high affinity binding partner of FGF2 provides insights explaining its prominent role in driving unconventional secretion of FGF2 from cells.

Based on the results presented in this study, one may wonder whether GPC1 is also relevant for unconventionally secreted proteins other than FGF2. While a comprehensive analysis looking into this aspect will be a goal of future studies, we tested galectin-1 and galectin-3 as potential binding partners of GPC1. They belong to a family of lectins that are known to be secreted by unconventional means (Rabouille, 2017; Dimou and Nickel, 2018; Popa et al., 2018). In contrast to FGF2, despite being lectins as well, both galectin-1 and galectin-3 were incapable of interacting with GPC1. These findings suggest that GPC1 is not a general component of unconventional secretory processes but may represent a highly specific molecular component driving FGF2 secretion.

Another intriguing finding of this study was the observation that GPC1 is dispensable for FGF signaling. This was evident from experiments demonstrating that FGF2-induced signaling leads to similar levels of phosphorylated ERK1/2 levels in GPC1 knockout versus wild-type cells. Likewise, overexpression of GPC1 did not affect FGF2-induced activation of ERK1/2. These findings suggest that other heparan sulfate proteoglycans such as syndecans are sufficient to support FGF2 signaling. By contrast, unconventional secretion of FGF2 largely depends on the presence of GPC1 as the key factor that determines the efficiency of this process. With these observations, while GPC1 is not essential for FGF signaling, our study reveals an intimate relationship between FGF2 and GPC1 in the secretion of FGF2 from cells. Since both FGF2 and GPC1 are key components of tumor progression for a wide range of cancer types (Akl et al., 2016; Pan and Ho, 2021), we propose that the prominent function of GPC1 in driving efficient unconventional secretion of FGF2 might play a key role for tumor development such as acute myeloid leukemia (Traer et al., 2016; Javidi-Sharifi et al., 2019).

## Acknowledgements

This work was supported by grants from the Deutsche Forschungsgemeinschaft (WN; SFB/TRR 83 and SFB/TRR 186). We are indebted to Julien Bethune (HAW Hamburg, Germany) who helped designing the BioID screen reported in this study. We would like to further acknowledge help from Holger Lorenz (ZMBH imaging facility) and Monika Langlotz (ZMBH FACS facility). We thank Marco Binder (DKFZ Heidelberg) for providing HEK293 cells used for the production of recombinant heparan sulfate proteoglycans and the FingerPrints mass spectrometry unit at Dundee University contributing the proteomics analysis contained in this study.

## Materials and Methods

### Cell culture

Hela S3 cells were cultured in DMEM, supplemented with 10 % FCS and 100 IU/ml penicillin and 100 μg/ml streptomycin at 37°C with 95 % humidity and 5 % CO_2_. Human embryonic kidney EcoPack 2–293 cells (Clontech) were cultivated on collagen-coated (Collagen R; Serva Electrophoresis) plates under the same conditions. HEK293 cells were cultured under the same conditions. For protein purification, HEK293 cells were grown in EX-CELL® ACF CHO Medium (Sigma-Aldrich, C5467) supplemented with 100 IU/ml penicillin and 100 μg/ml streptomycin at 37°C with 95 % humidity and 5 % CO_2_. CHO K1 cells were cultured in α-MEM medium supplemented with 10% FCS, 2 mM glutamine, 100 U/ml penicillin and 100 μg/ml streptomycin at 37 °C with 95% humidity and 5% CO_2_.

### Generation of stable cell lines

For all experiments, stable cell lines were generated with a retroviral transduction system based on Moloney Murine Leukemia Virus as previously described (Engling et al., 2002). Virus production was performed in HEK293 cells with a stably integrated pVPack-Eco packaging system in its genome as well as the retroviral packaging proteins (EcoPack 2–293 cells). Proteins expressed upon induction with doxycycline, like FGF2-GFP were cloned into the pRevTre2 vector, containing a Tet-response element and GPCs and SDCs were cloned into the pFB NEO vector. Retrovirus production was performed according to the MBS Mammalian Transfection Kit (Agilent Technologies) and virus was harvested after two days from confluent cells. Hela S3 and CHO K1 cells constitutively expressing the murine cationic amino acid transporter MCAT-1 (Albritton et al., 1989) and a Tet-On transactivator, rtTA2-M2 (Urlinger et al., 2000) were transduced with the freshly harvested virus. GFP expressing cells were selected by FACS, untagged protein containing cells were selected with G418 and protein expression levels were analyzed by Western blot. Heparan sulfate proteoglycans were detected after Heparinase III digest (NEB) using a monoclonal antibody (3G10) direct against the glycosylation attachment site in the core protein structure (370260, abcam).

### Generation of knock-out cell lines

Knockout cells were generated via CRISPR-Cas9 as previously described (Ran et al., 2013). Briefly, gRNAs for GPC1 (exon 2: fwd 5’-CACCGTGCAGCAGGTGTAGCCCTG-3’; rev 5’-AAACCAGGGCTACACCTGCTGCAC-3‘) and GPC5 (exon 3: fwd 5‘-CACCGATACTCAGAATGCATCCGGA-3‘; rev 5‘-AAACTCCGGATGCATTCTGAGTATC-3‘) were subcloned into pSpCas9(BB)-2A-RFP [based on pSpCas9(BB)-2A-GFP (PX458); (Ran et al., 2013)] using BbsI (NEB #R3539). GPC1, GPC5 and GPC1/5 knock-outs were generated in HeLa S3 FGF2-GFP cells that were grown in 6-well plates to 80% confluency. Cells were transfected with 2 μg DNA using FuGENE® HD Transfection Reagent (REF E2311, Promega). After 24 hours, cells were transferred to a 10 cm dish and FGF2-GFP expression was induced via addition of doxycycline (1μM). 48 h after transfection single cell clones were sorted for GFP and RFP fluorescence into 96-well plates. Clones were validated for GPC1/5 knockouts via Western analysis following heparinase III digestion using a monoclonal antibody (3G10) direct against the glycosylation attachment site in the core protein structure (370260, abcam) or via sequencing (GPC1: 5’-ACTCACCATCGAAGCTG-3’ and GPC5: 5’-GCGGCTGGGCAGCAGGGACCT-3’) as indicated.

### Identification of proteins in proximity to FGF2 in cells

Cells were detached with Gibco® Cell Dissociation Buffer (Thermo Fisher Scientific) and equal cell numbers were lysed in 2 ml Cyto0.2 Buffer (40mM HEPES pH7.4, 120mM KCl, 2mM EGTA, 0.4% glycerol, 0.2% NP-40, protease inhibitors) for 30min at 4°C while rotating. Nuclear fraction was pelleted for 3min at 1000xg, 4°C and the cytosolic fraction was supplemented with 0.4% SDS. After washing the pellet with Cyto0.1 Buffer (40mM HEPES pH7.4, 120mM KCl, 2mM EGTA, 0.4% glycerol, 0.1% NP-40, protease inhibitors) and Cyto0.0 Buffer (40mM HEPES pH7.4, 120mM KCl, 2mM EGTA, 0.4% glycerol, 0.0% NP-40, protease inhibitors), the nuclear fraction was lysed with Lysis Buffer (50mM Tris pH7.4, 500mM NaCl, 0.4% SDS, 5mM EDTA). Both fractions were sonicated four times for 30s at 4°C and centrifuged for 10min at 16.000xg to remove debris.

Hela S3 cells stably expressing myc-tagged BirA* or myc-tagged FGF2-BirA* under a doxycycline-inducible promoter were supplemented with doxycycline (1μM) for 48h and biotin (50μM) during the last 36h of culture. Cytosolic fractions generated as described above were adjusted to 2% Triton-X-100 and 150mM NaCl using 50mM Tris pH7.4 before incubation with 200μl streptavidin-coupled Dynabeads (M-280, 6.7x 108 beads/ml, Invitrogen) overnight at 4°C while rotating. Beads were washed two times for 8 min while rotating with each buffer: W1 (2% SDS); W2 (0.1% sodium deoxycholate, 1% TritonX-100, 500mM NaCl, 1mM EDTA, 50mM HEPES, pH 7.4); W3 (10mM Tris pH8, 250mM LiCl, 1mM EDTA, 0.5% NP-40, 0.5% sodium deoxycholate) and W4 (50mM Tris pH7.4, 50mM NaCl, 0.1% NP-40). Proteins were eluted in 4x sample buffer (40% glycerol, 240mM Tris-HCL pH6.8, 8% SDS, 5% β-mercaptoethanol and bromophenol blue) for 15 min at 95°C and separated in a 1.5mm 10-well 4-12% pre-cast gradient gel. After gels were washed three times with ddH20 for 10 min, a Colloidal Coomassie staining (0.02% CBBG-250, 5% aluminum sulfate (14,18)-hydrate, 10% EtOH, 6.8% orthophosphoric acid) was performed overnight at RT and afterwards destained (10% EtOH, 1.7% orthophosphoric acid) for 1 h. Gels were washed twice with ddH20 for 10 min before bands were cut and sent to FingerPrints Proteomics (Dundee University, Scotland).

Gel pieces were subjected to in-gel reductive alkylation and trypsin digest. Peptides were separated using strong cation exchange (SCX) fractionation before 1D nano-LC-MS/MS of each SCX fraction. The resultant mass spectrometry data from each fraction was merged prior to Mascot database search using the MaxQuant software (V1.5.5.1) with the human Uniprot sequence database (UP000005640, 9606 – Homo sapiens, 26.08.2015) for protein identification. Digestion mode was set to specific Trypsin/P with maximal 2 missed cleavages. Carbamidomethyl was set as fixed modification, acetylation of the N-terminus, oxidation (M) and biotinylation at lysine were chosen as variable modifications. Protein quantification was performed with unique and razor peptides. Protein intensities from the resulting ProteinGroups.txt file of three independent biological replicates were analyzed using the Perseus software (V1.5.5.1). Only proteins detected in the FGF2-BirA sample of at least 2 out of 3 biological replicates were considered for the analysis. The fold change between the FGF2-BirA and BirA-group was calculated from the means and log2-tansformed with standard imputation based on normal distribution. Significant differences between the groups were analyzed by 2-sided t-test.

### FGF2 secretion experiments based on cell surface biotinylation

3 × 10^5^ cells were seeded 48 h prior to biotinylation and incubated with 1 μg/ml doxycycline after 24 h for induction of FGF2-GFP expression. For biotinylation cells were placed on ice and washed twice with PBS-Ca/Mg (1 mM MgCl_2_, 0.1 mM CaCl_2_). Cells were incubated with 1 mg/ml sulfo-NHS-SS-biotin (Thermo Fischer Scientific, 21331) in incubation buffer (150 mM NaCl, 10 mM triethanolamine pH 9.0 and 2 mM CaCl_2_) on ice for 30 min with shaking, subsequently washed once with quenching buffer (100 mM glycine in PBS-Ca/Mg) and quenched for 20 min while shaking. Cells were washed twice with PBS and lysed for 10 min in lysis buffer (62.5 mM EDTA pH 8.0, 50 mM Tris-HCL pH 7.5, 0.4% sodium deoxycholate, 1% NP40 and protease inhibitors from Roche) at 37°C. Lysed cells were detached via scraping and transferred into an Eppendorf tube. Cells were sonicated 3 min in a sonification bath and incubated 15 min at room temperature with vortexing every 5 min to solubilize all proteins. Lysates were cleared via 10 min centrifugation at 13000rpm 4°C in a table top centrifuge. Meanwhile, Pierce™ Streptavidin UltraLink™ Resin (Thermo Fischer Scientific, 53114) was washed twice in lysis buffer via 1 min centrifugation at 3000 g. 5% input was taken from cleared lysates, mixed 1:1 with 4X sample buffer (40% glycerol, 240 mM Tris-HCL pH 6.8, 8% SDS, 5% β-mercaptoethanol and bromphenol blue) and boiled for 10 min at 95°C. The remaining lysate was incubated for 1 h at room temperature with over-head turning. Beads were spun down and washed once with wash buffer 1 (0.5 M NaCl in lysis buffer) and thrice with wash buffer 2 (0.5 M NaCl in lysis buffer containing 0.1% NP-40) via centrifugation. Beads were eluted via boiling in 4X sample buffer at 95°C for 10 min.

### FGF2 secretion experiments based on flow cytometry

After induction with Doxycycline (1 μg/ml) for 16 h in 6-well plates, cells were washed once with PBS and collected after treatment with PBS supplemented with 5 mM EDTA. Cell surfaces were stained with 300 μl complete medium containing anti-FGF2 antibody [1:100, (Engling et al., 2002; Zehe et al., 2006)] at 4 °C for 1 h. After centrifugation for 10 min at 500 g, pellets were washed with PBS and resuspended in 100 μl complete medium containing anti-rabbit APC antibody (1:500) followed by incubation for 30 min in the shaker at 800 rpm. After a final wash with PBS cells were recovered in 300 μl PBS for FACSCalibur™ Flow Cytometer (Becton Dickinson) measurement for detection of cell surface bound FGF2-GFP.

### Single Particle TIRF Translocation Assay

Quantification of secreted FGF2-GFP particles was achieved employing a previously established single particle TIRF assay (Dimou et al. 2019). Widefield fluorescence and TIRF images were acquired using an Olympus IX81 xCellence TIRF microscope equipped with an Olympus PLAPO ×100/1.45 Oil DIC objective lens and a Hamamatsu ImagEM Enhanced (C9100-13) camera. Data were recorded and exported in Tagged Image File Format (TIFF) and analyzed via Fiji (Schindelin et al., 2012). For the quantification of FGF2-GFP translocation to cell surfaces, CHO K1 cells were seeded in μ-Slide 8 Well Glass Bottom plates (ibidi) followed by incubation for 24 h in the presence of 1 μg/ml doxycycline to induce FGF2-GFP expression (for the experimental condition at high FGF2-GFP expression levels), or without doxycycline incubation (for the experimental condition at low FGF2-GFP expression levels). Following incubation, the medium was removed and cells were rinsed three times with Live Cell Imaging Solution (Thermo Fisher Scientific). Cells were further incubated on ice with membrane impermeable Alexa Fluor 647– labelled anti-GFP nanobodies (Chromotek) for 30 min. Afterwards, they were rinsed three times with PBS and fixed with 4 % PFA (Electron Microscopy Sciences) for 20 min at room temperature. GFP fluorescence was excited with an Olympus 488 nm, 100 mW diode laser. Nanobody fluorescence was excited with an Olympus 640 nm, 140 mW diode laser. The quantification of FGF2-GFP particles on cell surfaces was achieved through a quantitative analysis of TIRF images. The frame of each cell was selected by wide-field imaging. For the experimental condition at low FGF2-GFP expression levels, the EM Gain for the wide-field (GFP) was adjusted in order to properly select the cell area. The number of nanobody particles were normalized to the cell surface area (μm^2^). The total number of nanobody particles per cell was quantified employing the Fiji plugin TrackMate (Tinevez et al., 2017). Background fluorescence was subtracted for all representative images shown.

### Quantification of FGF2 binding to cell surfaces

Cells grown to confluency were washed with PBS and detached by cell dissociation buffer. After cell counting, 2 × 10^5^ cells were collected and washed again with PBS. Cell pellets were resuspended in 200 μl PBS and mixed with 200 μl of FGF2-GFP (5μg) followed by 1 hour incubation on a rotating wheel at RT. Cells were washed once with PBS and the pellet was resuspended in 200 μl PBS before analysis using a FACSCalibur™ flow cytometer (Becton Dickinson) for GFP intensities. Intensity values were normalised to WT cell intensities.

### Quantification of cellular glycosaminoglycan chains

Assays were performed according to the manufacturer’s Blyscan™ (biocolor) protocol. Briefly, cells were grown to confluency for 72 hours. After a PBS wash, cells were detached with PBS supplemented with 5 mM EDTA, and cells were dissolved in 400 μl Papain extraction buffer to be afterwards incubated for 6 h at 65 °C on a shaker. 100 μl of sample were mixed with 1 ml Blyscan dye reagent for 30 min while shaking and precipitated GAGs were pelleted for 10 min at 12.000 rpm in a centrifuge. After removal of supernatant, the pellet was dissolved in 500 μl dissociation reagent for 10 min in shaker and 200 μl of supernatant were analysed in 95 well plate at plate reader at 656 nm. Heparan sulfate chains were additionally analysed by incubation of 100 μl of papain digested GAG sample with sodium nitrite (100 μl), followed by addition of 100 μl acetic acid. After vortexing, the samples were incubated for 60 min at RT and nitrous acid was removed by addition of 100 μl ammonium sulphamate reagent for 10 min. 100 μl of neutralized sample were analysed as described above to quantify O-sulfated GAGS. Finally, to analyse the N-sulfated GAGs, results were subtracted from the total GAG amounts.

### Protein expression and purification

His-FGF2 (vector pQE30) and His-FGF2-GFP (vector pET15b) were purified from the E. coli strains W3110Z1 and BL21 Star, respectively. Following o/n expression at 25 °C, proteins were purified sequentially by Ni-NTA affinity chromatography (HisTrap FF, GE Healthcare), heparin chromatography (HiTrap Heparin HP, GE Healthcare) and size exclusion chromatography using a Superdex 75 column. Proteins were snap frozen in aliquots and stored at −80 °C. The GPCs indicated and SDC4 were cloned into the pcDNA3.1 vector containing a BM40 signal peptide (replacing of the original one) and a His-tag instead of a GPI anchor or a transmembrane domain (Fig. S2). Proteins were expressed in HEK293 cells and supernatants were harvested after four days. Following centrifugation and filtering (0.2 μm), proteins were purified via Ni-NTA affinity chromatography followed by size exclusion chromatography using a Superose 6 and Superdex 200 column.

### Biolayer interferometry to quantify protein-protein interactions

The biolayer interferometry allows a label-free analysis of real-time interaction events due to an optical detection of biomolecules that bind to the fiber-optic biosensors. Upon immobilization of the ligand to the biosensor a shift in the interference spectrum of the reflected light is induced and can be detected. As soon as the analyte binds the ligand, a further increase of the optical thickness on the biosensor surface is detected by the additional wavelength shift, which is then reported as the wavelength change (nm) over time (s). Measurements were performed on the OctetRed96e system (Sartorius) using the Streptavidin sensors (18-5019 SA, Sartorius). Data were evaluated with the Data Analysis HT 12.0 software (Sartorius). Proteins were measured in black 96-well plates (655209, Greiner) in 200 μl for all samples. Biosensors were hydrated for 10 min before measurements in Octet Buffer (PBS, 0.02% Tween and 0.1% BSA) to remove sucrose coverage. Measurements were conducted with the plates shaking at 1000 rpm. All assays were performed with SA biosensors and the ligands were biotinylated. Proteins were labeled with EZ-Link™ NHS-PEG4-Biotin (Thermo Scientific™ A39259) in a 1:1 ration at 37 °C for 30 min. Proteins were separated from non-bound biotin by Zeba™ Spin Desalting Columns (Thermo Scientific™ 89882). Biotinylation does not interfere with the binding kinetics, as the interaction takes place at the HS chains. A loading scout was performed to find the optimal amount of bound ligand to the biosensor surface. All kinetic experiments were performed with 6 μg/ml of biotinylated ligand loaded to the sensor for 10 min. If not stated otherwise in the figure legends, the assay set up was as follows: 1) baseline in Octet Buffer (2 min), 2) load with ligand (6μg/ml, 10 min), 3) wash in octet buffer (1 min), 4) baseline II in octet buffer (1 min), 5) association of FGF2 (60 μM dilution series, 1 or 5 min), 6) dissociation in octet buffer (1 or 5 min), 7) recovery in glycine pH 1.7 and octet buffer (3× 5 sec each).

### FGF2 signaling assays

Hela S3 cells with a FGF2 and GPC1 KO background were induced with recombinant FGF2 (1 and 10 ng/ml) for 20 min. In parallel, Heparinase digests were performed for 2 hours at 37 °C with a mixture of Heparinase I + II and III (3.5 mUnits/ml, NEB) in Heparinase digestion buffer (20 mM Tris, 100 mM NaCl and 1.5 mM CaCl_2_, pH 7.4) before cell signaling was induced by FGF2. Cells were lysed and analyzed by Western Blot for ERK1/2 (4696, CST) and pERK1/2 (9101, CST) levels. GAPDH (AM4300, Invitrogen) was detected as loading control.

### Identification and quantification of heparan sulfate disaccharides

HS disaccharides of HSPGs were analyzed as described previously (Carnachan and Hinkley, 2017). Briefly, proteins (1mg/ml) were digested in 200 μl digestion buffer (100 mM NaOAc, 2 mM CaOAc, pH 7.0) with 1.75 mIU of Heparinase I + II and III (NEB) for 16 h at 30 °C. Heparinases were inactivated at 95 °C for 10 min and denatured proteins were pelleted at 16.000. g for 10 min. 100 μl of supernatant was analyzed by HPLC with a strong anion exchange column (ProPac™ PA1, Thermo Fisher) with an elution gradient of 2 M NaCl, pH 3.5. Disaccharides were detected at an absorbance maximum of 232 nm. A standard mixture of HS disaccharides (20 μg/ml; Iduron, UK) was analyzed to identify HS disaccharides.

## Figures and legends

**Fig. S1:**
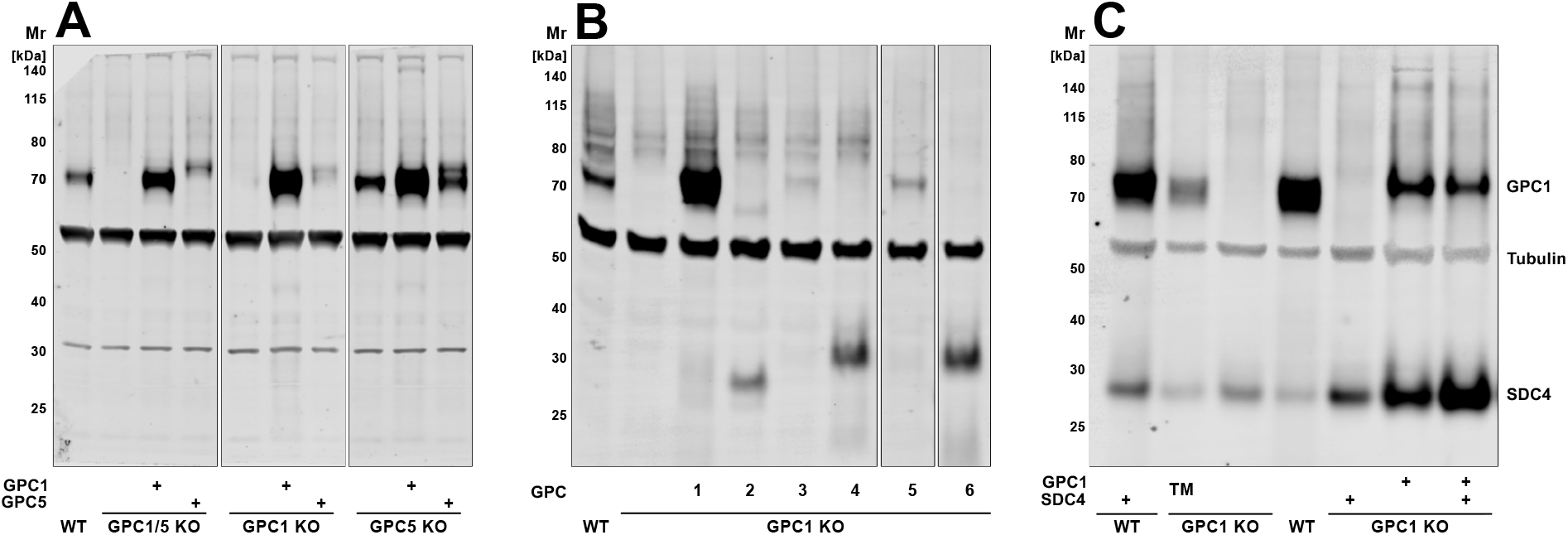
Characterization of engineered HeLa S3 cells used to quantify FGF2 secretion efficiencies. (A) HeLa S3 cell lines with different knockout backgrounds along with overexpression of GPC1 and GPC5, respectively, as indicated. (B) HeLa S3 cell lines with a GPC1 knockout background along with overexpression of GPC family members as indicated. (C) HeLa S3 cell lines with a GPC1 knockout background along with overexpression of either GPC1, SDC4 or a GPC1 construct in which the GPI anchor was replaced by a transmembrane span derived from SDC4. All panels represent Western analyses following digestion with heparinase III using antibodies directed against the heparan sulfate attachment site (3G10) to detect the proteins indicated. For experimental details, see Materials and Methods.

**Fig. S2:**
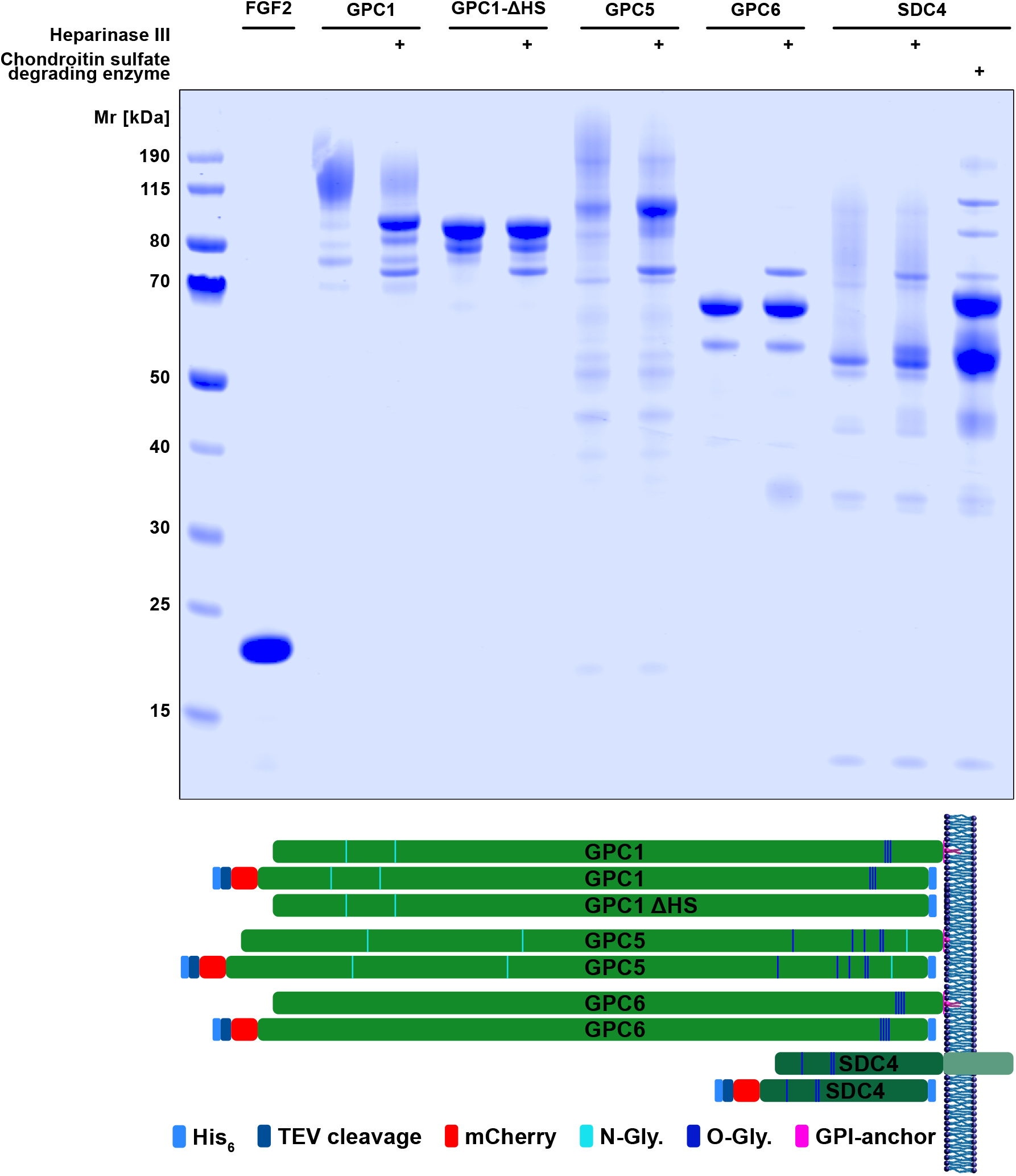
Expression and purification of soluble recombinant forms of FGF2 and the various heparan sulfate proteoglycans. FGF2 was expressed and purified from *E. coli* cells. All heparan sulfate proteoglycans indicated were expressed and purified from HEK293 cells. For details, see Materials and Methods. Upper panel: SDS-PAGE analysis of all purified proteins (Coomassie staining) Lower panel: Schematic description of constructs used in this study in comparison to the endogenous forms

**Fig. S3:**
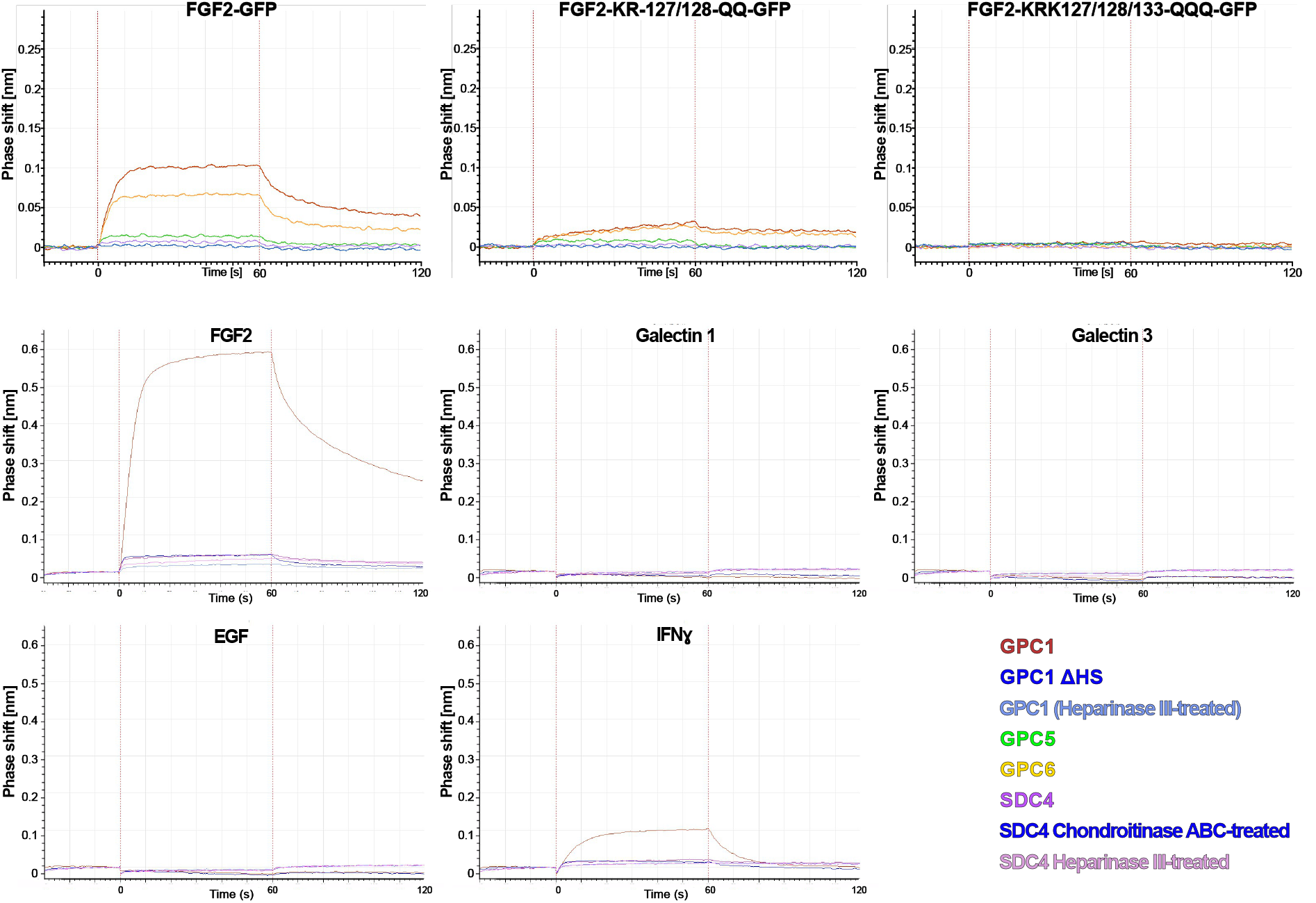
Interaction studies between different kinds of heparan sulfate proteoglycans and growth factors or cytokines using biolayer interferometry. Heparan sulfate proteoglycans were loaded onto BLI sensors as indicated by the color code. In some cases, they were treated with heparinase III or chondroitinase ABC as indicated. The binding partners were used in solution at a concentration of 60 nM. The data shown are representative for two independent experiments. For details, see Materials and Methods.

**Fig. S4:**
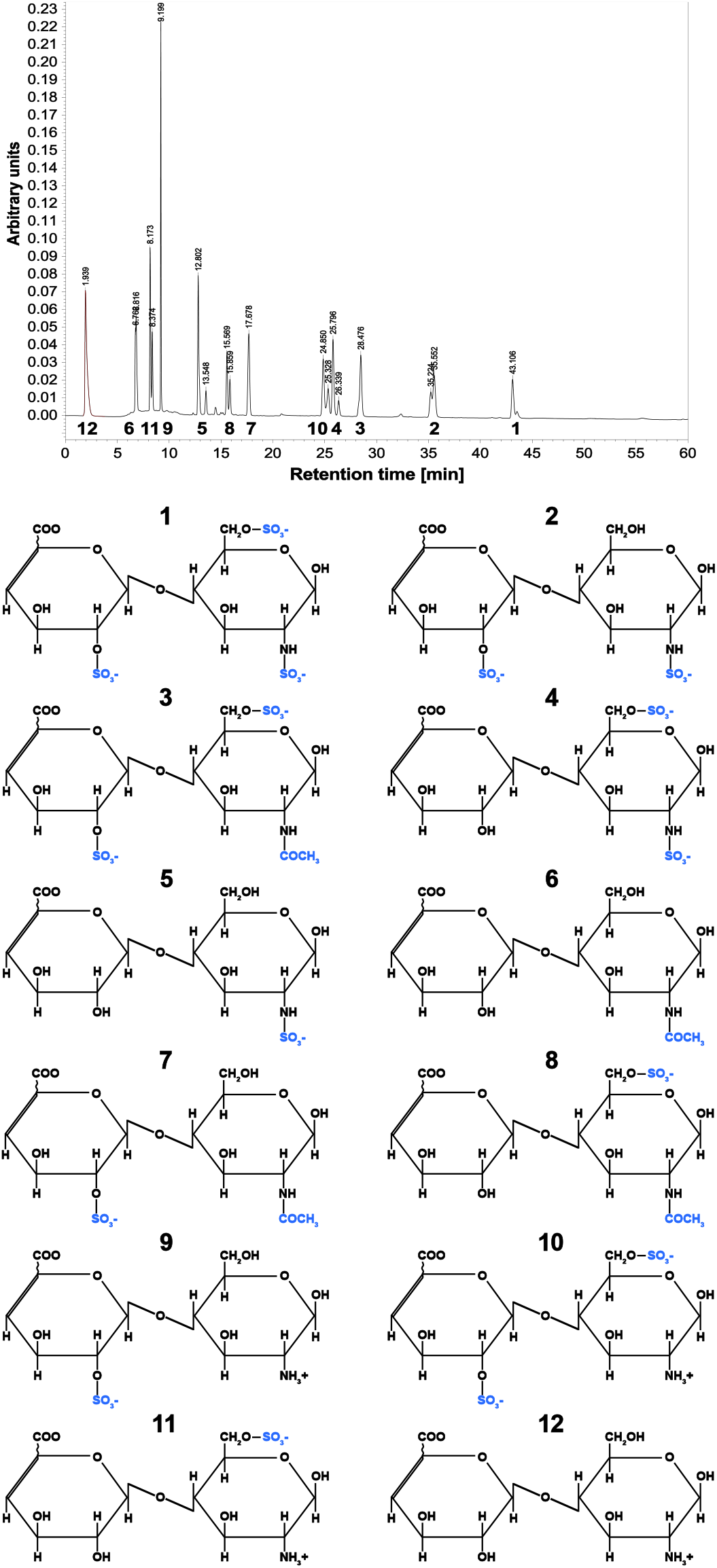
Characterization of synthetic dissacharide standards corresponding to the building blocks of heparan sulfate chains using an analytical HPLC analysis. The standards used were commercial products (Iduron, UK). They were characterized for their retention times on a strong anion exchange column operated by a HPLC system. For experimental details, see Materials and Methods

